# Leveraging single-cell transcriptomics of developing rat ocular outflow tissues to prioritize congenital glaucoma candidate genes

**DOI:** 10.1101/2025.08.26.672439

**Authors:** Sean M. Martin, Kristina N. Whisenhunt, Stuart W. Tompson

**Author notes:** Corresponding author (ST). These authors contributed equally to this work (Sean M. Martin, Kristina N. Whisenhunt). Commercial relationships disclosures: The authors declare no competing interest.

## Abstract

Primary congenital glaucoma (PCG) is a severe, early-onset eye disease most often caused by abnormal development of the aqueous humor outflow pathway (AHOP). Located circumferentially at the anterior chamber angle where the iris meets the cornea, the AHOP comprises the trabecular meshwork (TM) and Schlemm’s canal (SC), which together regulate intraocular fluid drainage. Malformation of this pathway leads to elevated intraocular pressure, painful ocular enlargement, and retinal damage that can result in blindness. While single-gene mutations account for approximately 25% of PCG cases across diverse populations, most molecular causes remain unknown and are likely due to rare or complex genetic factors. Progress in identifying these mechanisms has been limited by a lack of detailed gene expression data during AHOP development. To address this gap, we generated the first single-cell RNA sequencing dataset from developing AHOP tissue, using rat eyes at three key stages of TM and SC formation. This high-resolution dataset contains the transcriptomic profiles of 29,626 genes across 86,653 cells clustered into 13 general cell types, which included over 10,000 cells related to TM/SC subtypes. Analysis of 44 genes previously linked to Mendelian childhood glaucoma showed that 36 (82%) were expressed in these TM/SC-related populations, validating the dataset’s relevance. Notably, this study identified 395 genes selectively upregulated in developing TM/SC subtypes, revealing numerous candidates potentially involved in the formation and function of TM/SC structures. This resource will support the discovery of rare Mendelian disease genes and inform the development of polygenic risk scores for complex genetics underlying early-onset forms of glaucoma.

## Introduction

In the eye, aqueous humor (AH) is a clear fluid produced by the ciliary body that flows through the anterior chamber, delivering nutrients, clearing metabolic waste, and helping maintain ocular shape for focused vision. AH production is balanced via drainage through structures located at the interface between the cornea and iris – the conventional AH outflow pathway (AHOP). To prevent excess intraocular pressure (IOP), fluid exits the eye through the trabecular meshwork (TM) and Schlemm’s canal (SC), which connects to the systemic blood circulatory system via collector channels (Fig 1). Failure to correctly develop these intricate outflow structures can lead to elevated eye pressure, damage to retinal ganglion cells and the optic nerve, and early-onset vision loss known as primary congenital glaucoma.

**Fig 1.**
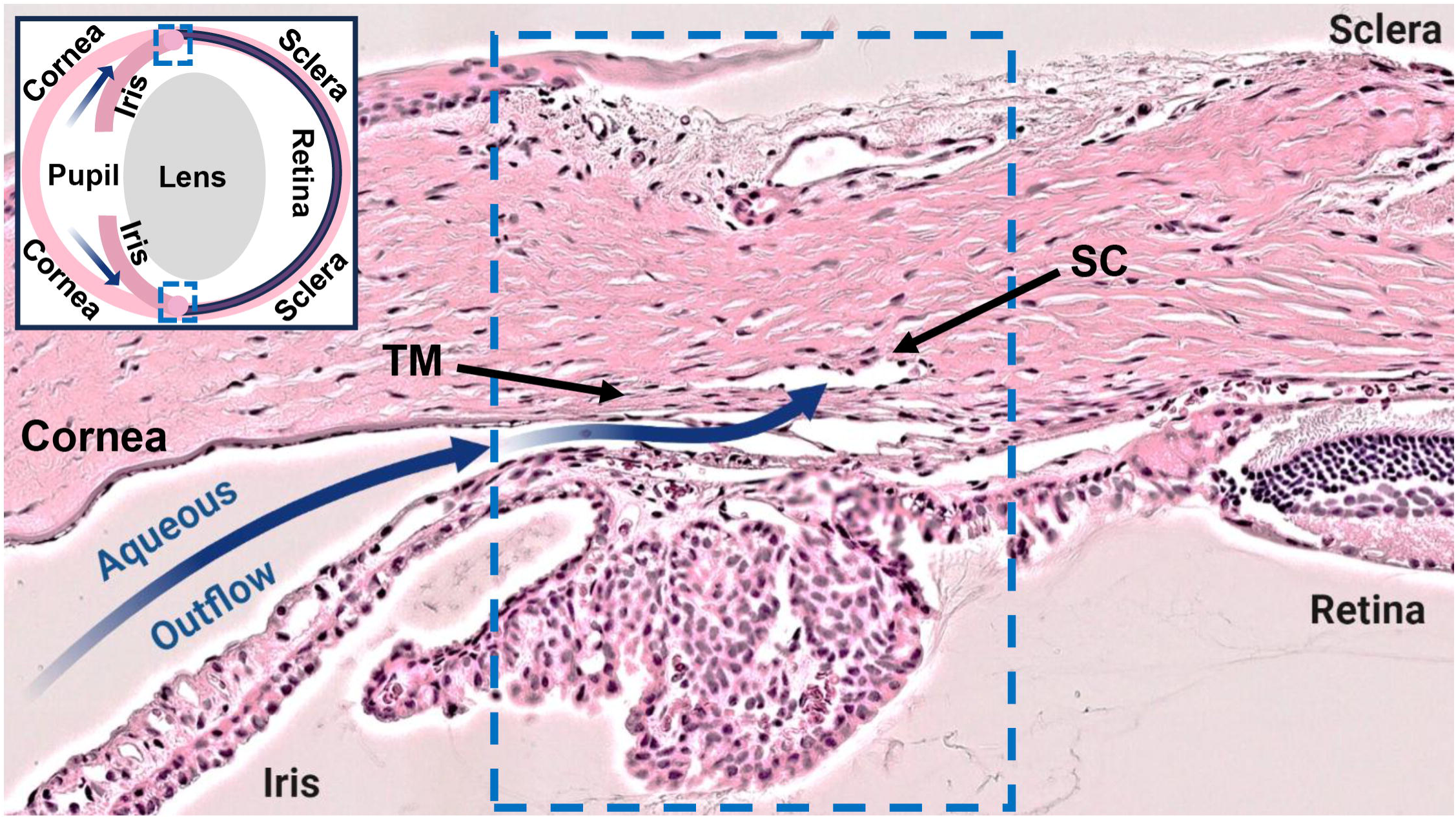
Route of aqueous humor outflow from the eye. H&E-stained tissue section from a mature (P80) rat eye showing the location of the AHOP at the junction of the cornea and iris. Aqueous humor exits the eye (blue arrows) through the trabecular meshwork (TM) and Schlemm’s canal (SC), which encircle the anterior chamber. Tissues dissected for scRNAseq profiling are highlighted (blue dashed box) and shown relative to a whole eye (insert).

The molecular mechanisms of primary congenital glaucoma have been elucidated in only 25% of cases, and all involved genes critical for the proper formation and function of the TM and/or SC tissues.^1–4^ Our understanding of genes important for AHOP development has been restricted by the limited availability and complex heterogeneity of these early tissues. Several studies have recently employed single-cell RNA sequencing (scRNAseq) to catalogue gene expression in outflow tissues from humans, nonhuman primates, pigs and mice,^5–9^ but all profiled mature tissues. To date, no study has attempted to identify the cells and molecular pathways active during the early development and growth of the TM and/or SC tissues relevant to the etiology of congenital glaucoma.

The mature TM is a sieve-like structure composed of three distinct layers, each contributing to the regulation of AH outflow and IOP.^10^ AH first passes through the uveal meshwork – a loose, open network of connective tissue encased in a continuous basement membrane and lined with endothelial- like TM cells (TMCs). This layer serves to filter larger cellular debris and neutralizes reactive oxygen species (ROS). The middle corneoscleral layer consists of TMC-covered collagen and elastin-rich beams with smaller pores for finer filtration and additional ROS scavenging. Finally, the juxtacanalicular tissue (JCT), composed of loosely arranged extracellular matrix (ECM) and star-shaped fibroblast-like TMCs, regulates AH outflow resistance in coordination with the inner wall of SC.^11^ The mature SC is composed of a monolayer of vascular endothelial cells that share molecular characteristics with blood and lymphatic endothelial cells.^12,13^ The SC endothelial cells (SECs) can be divided into inner-wall (IW) cells that lay against the TM and outer-wall (OW) cells located towards the outside of the eye. The IW allows AH outflow through giant vacuoles and pores whilst maintaining a blood-aqueous barrier, whereas the OW maintains the canal’s structure and passes AH into collector channels that lead away from the eye.^14^

Here, we present a study utilizing scRNAseq to identify the major cell types and their associated gene expression profiles during three major stages of SC and TM development in rats. Rat tissues share all major stages of anterior chamber angle development and morphological characteristics of the AHOP with humans, and they also enable acquisition of larger quantities of relevant tissue components versus mice. Remé and colleagues performed detailed electron and light microscopy studies of rat AHOP development,^15^ observing AHOP anlage formation at embryonic day 7. By postnatal day 5 (P5), the presumptive TM remained an undifferentiated condensation of periocular mesenchyme (POM), and a rudimentary cluster of blood vessels that would become SC were visible. Between P5 and P10, sheets of primitive TM formed, and the young SC vessels became enlarged. Between P10 and P20, the TM became five or six layers with distinct cells and SC had fused into a vessel with a single vacuole. The final structures were formed between P20 and P60, when the TM became less dense with the formation of Fontana’s spaces and beams, and both the TM and SC elongated into their mature forms. Based on these developmental observations, we profiled these specialized tissues at P7-9, P14-16, and P27-29.

## Results

### Limbal cell acquisition

Histological examination of rat AHOP tissues at P8, P15, and P28 confirmed the morphological findings described by Remé and colleagues (Fig 2).^15^ In total, three male and three female limbal tissue strips (region highlighted in Fig 1) were dissected and dissociated into single-cell suspensions at each of the three developmental ages. Samples contained all cell types located between the cornea and sclera on the exterior surface of the eye to the ciliary body on the inner side. The final dataset contained transcriptomic profiles of 29,626 genes from 86,653 limbal cells, visualized as 13 major cell type clusters (C0-C12) according to overall gene expression similarity (Fig 2). The contributions of cells from the 18 tissue samples to each cluster are provided in S1 Table. The major cell type identities within these clusters were revealed through differential expression of published marker genes, as described in Supporting Information (“Identification of major limbal cell types” in S1 Appendix and Fig S1).

**Fig 2.**
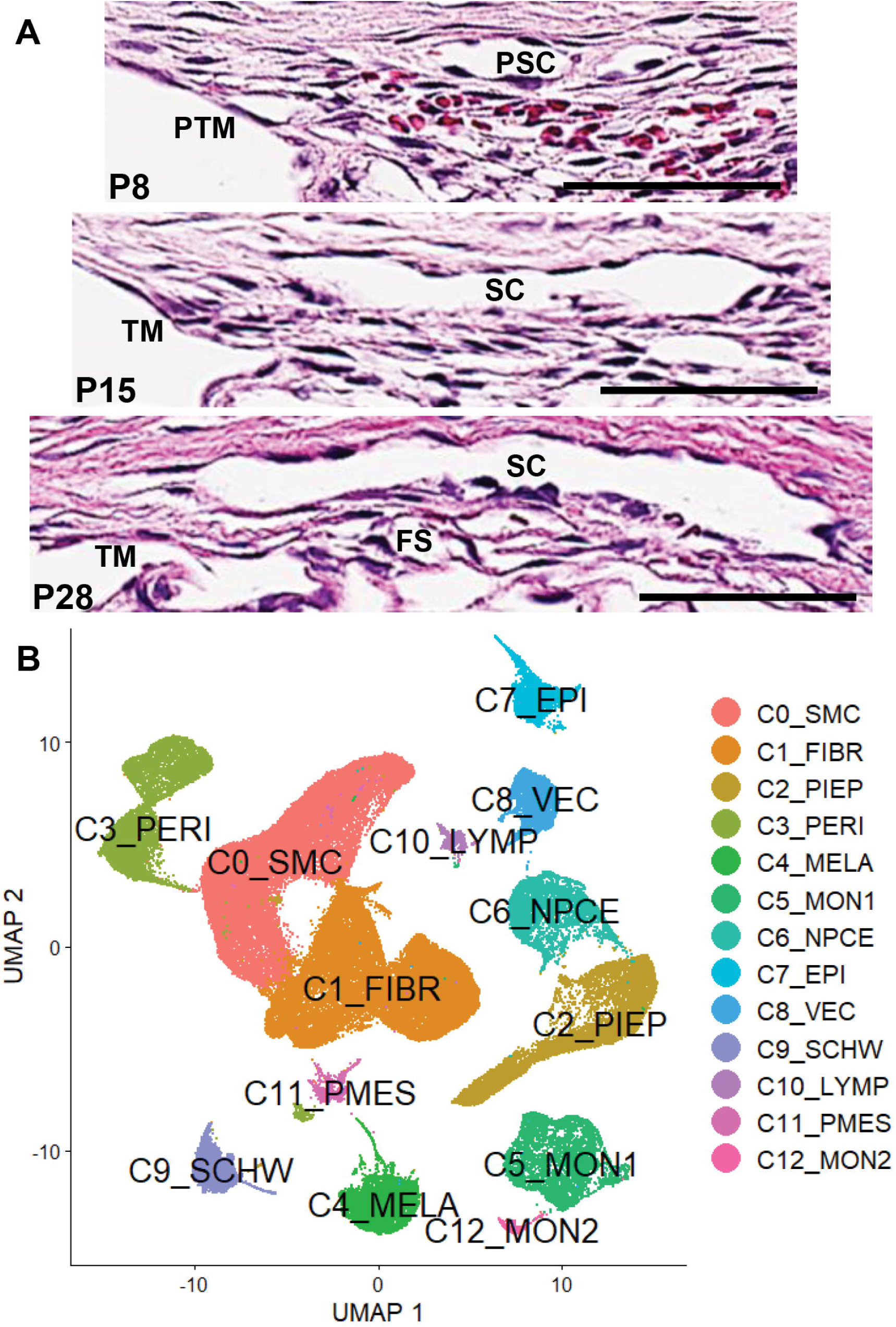
Morphology of dissected developing AHOP tissues and resulting scRNAseq clustering of complete limbal dataset. (A) H&E-stained anterior eye sections highlighting the AHOP tissues at P8, P15, and P28. Labels show primitive TM (PTM), primitive SC (PSC), trabecular meshwork (TM), Schlemm’s canal (SC), and Fontana’s spaces (FS). Scale bars 50µm. (B) Clustering of single-cell profiles visualized by two-dimensional Uniform Manifold Approximation and Projection (UMAP). The 13 major clusters are distinguished by different colors and numbered (C0-C12) by descending cell population. Cell type identities assigned by key marker gene expression (Fig S1) are smooth muscle cells (SMC), fibroblasts (FIBR), pigmented epithelium (PIEP), pericytes (PERI), melanocytes (MELA), monocytes-1 (MON1), nonpigmented ciliary epithelium (NPCE), surface epithelium (EPI), vescular endothelial cells (VEC), Schwann cells (SCHW), lymphocytes (LYMP), proliferating mesenchyme (PMES), and monocytes-2 (MON2).

### Vascular endothelium and the identification of Schlemm’s canal cells

Vascular endothelial cells (VECs), including those that form Schlemm’s canal (SECs), uniquely express *Pecam1* and formed a single independent cluster (C8, Fig S2). To specifically identify SECs, these 3,155 cells were reclustered into nine subtypes (VEC0-VEC8, Fig 3 and Fig S3). Their vascular identities were ascertained through interrogation of differentially expressed genes, as described in Supporting Information (“Identification of VEC subtypes” in S1 Appendix and Fig S3). In mice, SECs share transcriptional features of both blood and lymphatic endothelia, expressing *Cd34* and *Prox1*, but not *Pdpn* or *Vwf*, yet they also retain distinct gene expression patterns that reflect their specialized role in AH drainage.^9,12,13,16^ SECs expressing these hybrid features formed VEC1 and a subpopulation of VEC7 (Fig 3 and Fig S3). Consistent with the growth and maturation of SC, the SECs population (VEC1) dramatically increased with sample age (Fig 3 and Fig S4). The two distinct SEC-related populations in VEC7 were formed from proliferating cells highly expressing *Top2a* and *Mki67*.^17,18^ These populations also changed in size with tissue age (Fig S4). As SC developed, the larger population displaying a stronger blood vessel-like profile (high *Flt1*, *Lrg1*, *Stc1*, *Sema3e*, *Tcf15*, low *Prox1*) reduced in number, whilst the smaller island exhibiting more SEC-like expression became numerous. This was consistent with cells involved in the formation of SC - proliferating VECs shifting their expression profile during canalogenesis from blood vessel-like to a more hybrid-lymphatic phenotype and ultimately contributing to the more specialized SEC pool in VEC1.^12^

**Fig 3.**
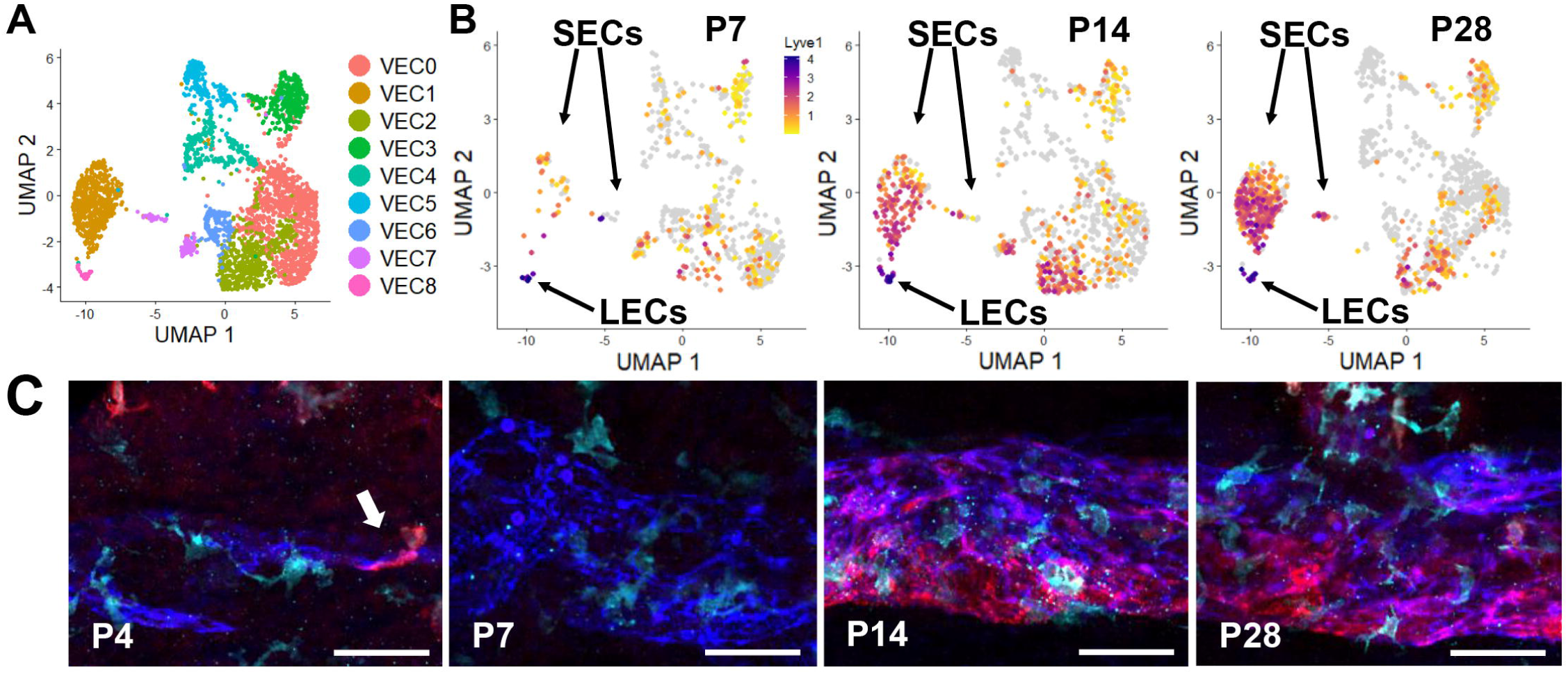
As SC develops in rats, a population of SECs express *Lyve1* as they take on a more lymphatic phenotype. (A) Reclustering of 3,155 VECs from the limbal C8 population visualized by UMAP in nine subtypes (VEC0-VEC8). (B) *Lyve1* expression throughout the VEC dataset (normalized log2 fold change expression). (C) Whole mount anterior cup tissues stained with Pecam1 (blue) to visualize SC, Aif1+ (cyan) macrophages, and Lyve1+ (red) subsets of macrophages and SECs. Macrophage-aided fusion of rudimentary vessels shown at P4 (arrow). Images oriented SC inner wall down. Scale bars 50µm.

In mouse studies, SECs do not significantly express the *Lyve1* gene, which is restricted to lymphatic vessels and a subset of interstitial macrophages that preferentially localize near vessels.^9,12,13,16,19^ In contrast, our studies in rat have revealed *Lyve1* expression not only in lymphatic vessels and interstitial macrophages but also at low levels in more mature SECs (Fig 3). We confirmed this finding at the protein level, immunostaining whole mounted tissues for Pecam1 to visualize SC, Aif1 to broadly highlight macrophages, and Lyve1 (Fig 3).^20^ At P4, areas of SC consisted of rudimentary smaller vessels that did not express Lyve1, and could be observed undergoing macrophage-aided vessel fusion (arrow). At P7, regions of SC comprised clustered vessels that were not fully fused and Lyve1 levels remained insignificant. At P14, Lyve1 expression by SECs was distinct from resident macrophages and was stronger towards the inner wall. At P28, Lyve1 expression by SECs was strongest and remained primarily towards the inner wall, reflecting the acquisition of lymphatic-like properties by the mature outflow vessel.

### Identification of SEC subtypes

In total, 655 VECs with SC-like profiles were identified, comprising 109 actively proliferating cells and 546 that were quiescent and more specialized. To resolve further SEC heterogeneity, VEC1 and VEC7 cells were isolated and re-clustered, yielding six subpopulations with distinct gene expression profiles (SEC0-SEC5, Fig 4; heatmap of top differentially expressed genes between SEC clusters provided in Fig S5). To support this dataset, *in situ* hybridization (ISH) studies were performed on AHOP tissues to confirm the localized expression of key SEC markers (Fig S6). The method used fluorescent probes to detect individual mRNA transcripts in and around cell nuclei in intact but thin tissue sections. Due to the limited number of nuclei captured from sections across the lumen of SC, this technique was only able to effectively localize robustly expressed genes. Furthermore, as observed for *Pecam1*, *Tek*, *Flt1*, and *Cd34*, transcripts were more readily detected in IW versus OW cells, despite their well-established expression throughout SC (Fig S6).

**Fig 4.**
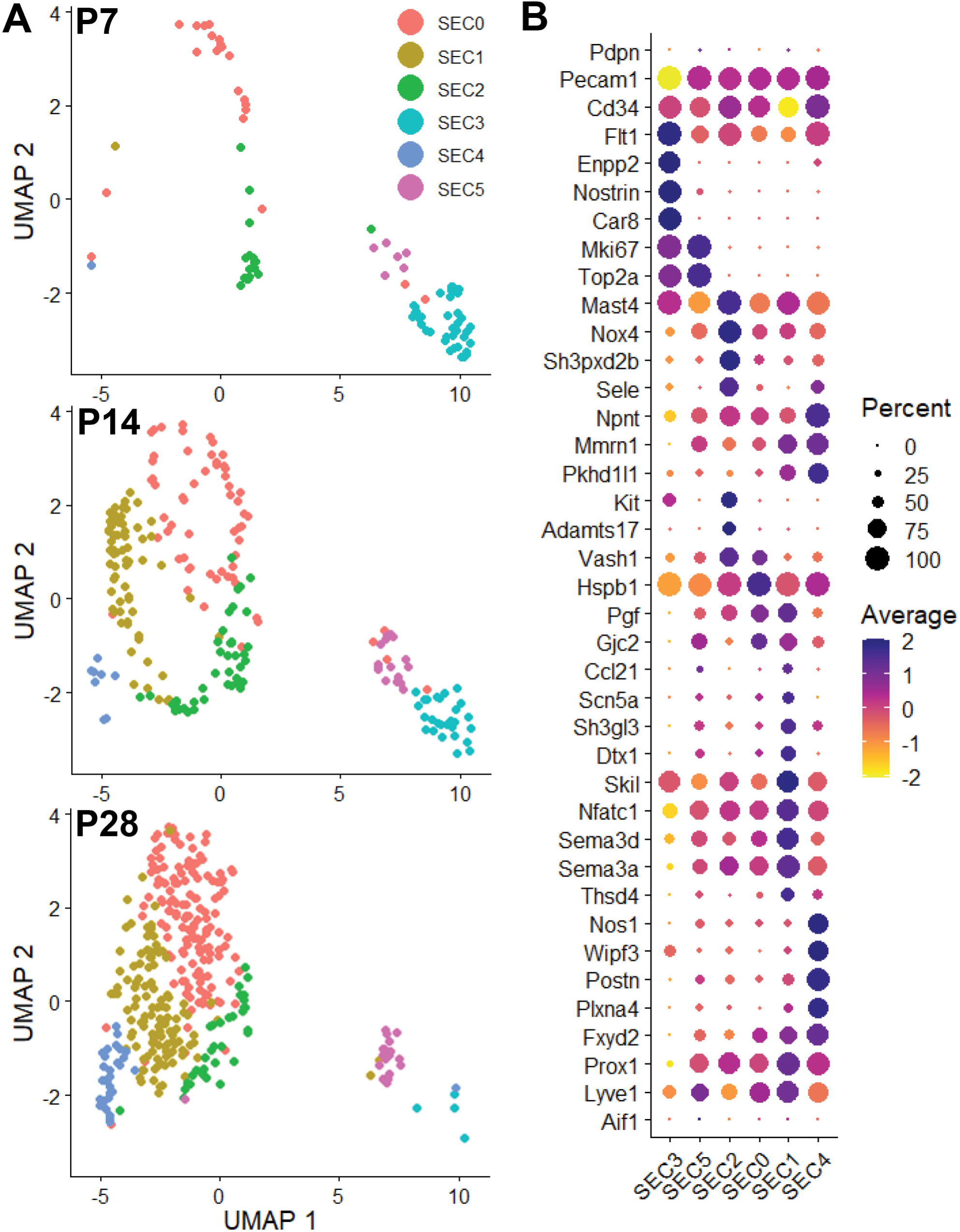
Cellular subtypes and associated gene expression within the Schlemm’s canal endothelial cell population during development. (A) UMAP plot of the six SEC subpopulations at the three developmental timepoints. (B) Dot plot of key marker gene expression distinguishing SEC subtypes. The dot diameter represents the percentage of each cell type expressing the gene, while the color denotes the scaled average expression within the cluster.

SECs derived from VEC1 formed four subclusters, while SECs from VEC7 remained in two spatially isolated populations (SEC3 and SEC5) distinguished by strong expression of genes related to cell replication. SEC3 cells were most abundant during early SC development (P7) and expressed genes associated with angiogenic sprouting and proliferation, including *Top2a*, *Mki67*, *Car8*, *Enpp2*, and *Nostrin*.^17,18,21–23^ In contrast, SEC5 cells lacked sprouting markers but expressed genes for lymphatic processes (*Prox1*, *Lyve1*, and *Nfatc1*,^24^) and vessel remodeling (*Sema3a*^25^, *Sema3d*^26^). ISH studies of *Top2a* expression were unable to localize SEC3/SEC5 cells to a specific SC region (Fig S6), which reflected their limited population at any age.

SEC2 cells were observed during mid-stage development of SC and displayed transcriptional features similar to SEC5 but without proliferation markers. Specifically upregulated expression was related to progenitor/stem cell identity (*Mast4*,^27^ *Kit*^28^), leukocyte chemotaxis (*Sele*,^29^), motility (*Sh3pxd2b*^30^), ECM modification (*Adamts17*^31^), and angiogenesis regulation (*Nox4*,^32^ *Vash1*^33^). The expression of *Sele* has previously been attributed to OW cells in adult mice.^9^ ISH studies of *Sele* expression in whole tissues reflected the scRNAseq data, with transcripts more readily detected later in SC development, but the signal was too limited to confirm localization to a specific region such as the OW (Fig S6). Conversely, *Nox4* expression signal was strong throughout AHOP development in both the IW and OW at P7 and P14 but appeared more restricted to the IW at P31 (Fig S6).

The largest late-stage populations of SEC0 and SEC1 cells showed closely related profiles, including expression of a gap junction protein associated with lymphatic-like endothelia, *Gjc2*.^34^ ISH studies of *Gjc2* localized the most robust expression to IW cells at P14 (Fig S6). Specifically, SEC0 cells showed the strongest expression of *Gjc2* and *Hspb1*, a protein involved in endothelial movement.^35^ Conversely, SEC1 cells exhibited enhanced expression related to lymphatic identity (*Prox1*, *Lyve1*, *Nfatc1*), vessel remodeling (*Sema3a*, *Sema3d*), regulation of angiogenesis (*Skil*^36^, *Pgf*^37^), ECM remodeling (*Thsd4*^38^), and a chemokine that attracts immune cells toward lymphatic-like vessels (*Ccl21*^39,40^). High *Prox1* and *Ccl21* expression have previously been associated with IW cells in adult mice.^9^ Consistent with this, ISH studies of *Ccl21* identified low expression localized to the IW (Fig S6). Furthermore, SEC1 cells also expressed genes that could be related to IW function, such as flow–responsive lymphatic remodeling (*Dtx1*^41,42^), lumen maintenance (*Sh3gl3*^43^), and sodium transport (*Scn5a*^44^). The smallest late-stage population of SEC4 cells were enriched for genes involved in integrin signaling (*Postn*^45^), cytoskeletal dynamics (*Wipf3*^46^), vascular integrity (*Plxna4*^47^), and maintenance of vascular homeostasis (*Nos1*^48^). Like SEC2, SEC4 cells also expressed *Sele*, which suggested they were derived from the OW. While enhanced expression of the ECM component nephronectin (*Npnt*^49^) has been reported for a subset of IW cells in adult mice,^9^ this gene was more broadly expressed in all rat SECs with particular enrichment in SEC4 (putative OW cells). Interestingly, both SEC1 (IW) and SEC4 (OW) cells shared expression of two genes previously identified in human lymphatics, *Mmrn1* and *Pkhd1l1*.^50^ Finally, most SEC clusters selectively expressed a subunit of the sodium/potassium ATPase pump that is critical for cell volume maintenance (*Fxyd2*).^51^ Given that *Fxyd2* expression in the rat limbus was otherwise restricted to monocytes, this gene emerged as one of the most specific markers for all SEC types in our dataset.

### Identification of trabecular meshwork cells within fibroblast-like populations

As demonstrated by Remé et al., in rats the TM begins forming as a condensed mass of multipotent periocular mesenchyme (POM) between P7-P9, which differentiates into primitive TM sheets.^15^ These early POM cells express high levels of the developmental transcription factor *Pitx2*,^52^ which is subsequently downregulated as they transition into TM cells (TMCs) - elongating, flattening, and becoming separated by extracellular fibers that organize into lamellae and connective tissue.^53^ By P14- P16 the layered TM structures are evident, and by P27-P29 the tissue resembles mature architecture with a meshwork of beams and large Fontana’s spaces.^15^

In the mature tissue, TMCs across all three layers express shared genes, including *Col4a1* and *Col8a1* (Type IV and VIII collagen alpha chains),^11^ *Eln* (elastin),^54^ *Nid1* (nidogen-1),^11^ *Myoc* (myocilin),^7^ *Mgp* and *Spp1* (calcification inhibitors),^55,56^ *Angpt1* and *Svep1* (angiogenic mediators),^8,57^ and *Ltbp2* (an ECM protein involved in TGF-β regulation).^58^ TMCs also express *Acta2* (smooth muscle actin),^59^ but not *Des* (desmin),^11^ distinguishing them from smooth muscle cells. Unlike the adjacent corneal endothelia, TMCs do not express *Col4a3*, encoding the α3 chain of type IV collagen.^11^

Layer-specific functions of TMCs are reflected by their differential gene expression. TMCs of the uveal and corneoscleral layers secrete tissue plasminogen activator (*Plat*) to maintain pathway patency,^60^ superoxide dismutases (*Sod1*, *Sod2*, *Sod3*) to combat reactive oxygen species (ROS),^61^ and *Il6* for IOP regulation.^62^ In primates, hyaluronic acid (HA) is present in the JCT ECM but it is not synthesized locally.^63^ Instead, HA is produced by *Has1*/*2*/*3* in uveal and corneoscleral TMCs^63,64^ and is carried into the JCT by AH flow.^59^ JCT-resident TMCs are identifiable by elevated expression of chondroadherin (*Chad*),^5^ tenascin-C (*Tnc*),^65^ angiopoietin-like 7 (*Angptl7*),^5–7^ chitinase-3-like protein 1 (*Chi3l1*),^5^ thrombospondin-1 (*Thbs1*),^66^ and crystallin alpha B (*Cryab*).^11,67^

To identify TMCs in the rat limbal dataset, cells were screened for high expression of the POM marker *Pitx2*, along with four genes previously localized to the TM: *Myoc*, *Angpt1*, *Svep1*, and *Ltbp2*.^7,8,58^ These genes showed variable expression across several limbal clusters but all were strongly enriched in C1 (Fig 5). Consequently, subsequent analyses were focused on this population of 21,735 fibroblast-like cells.

**Fig 5.**
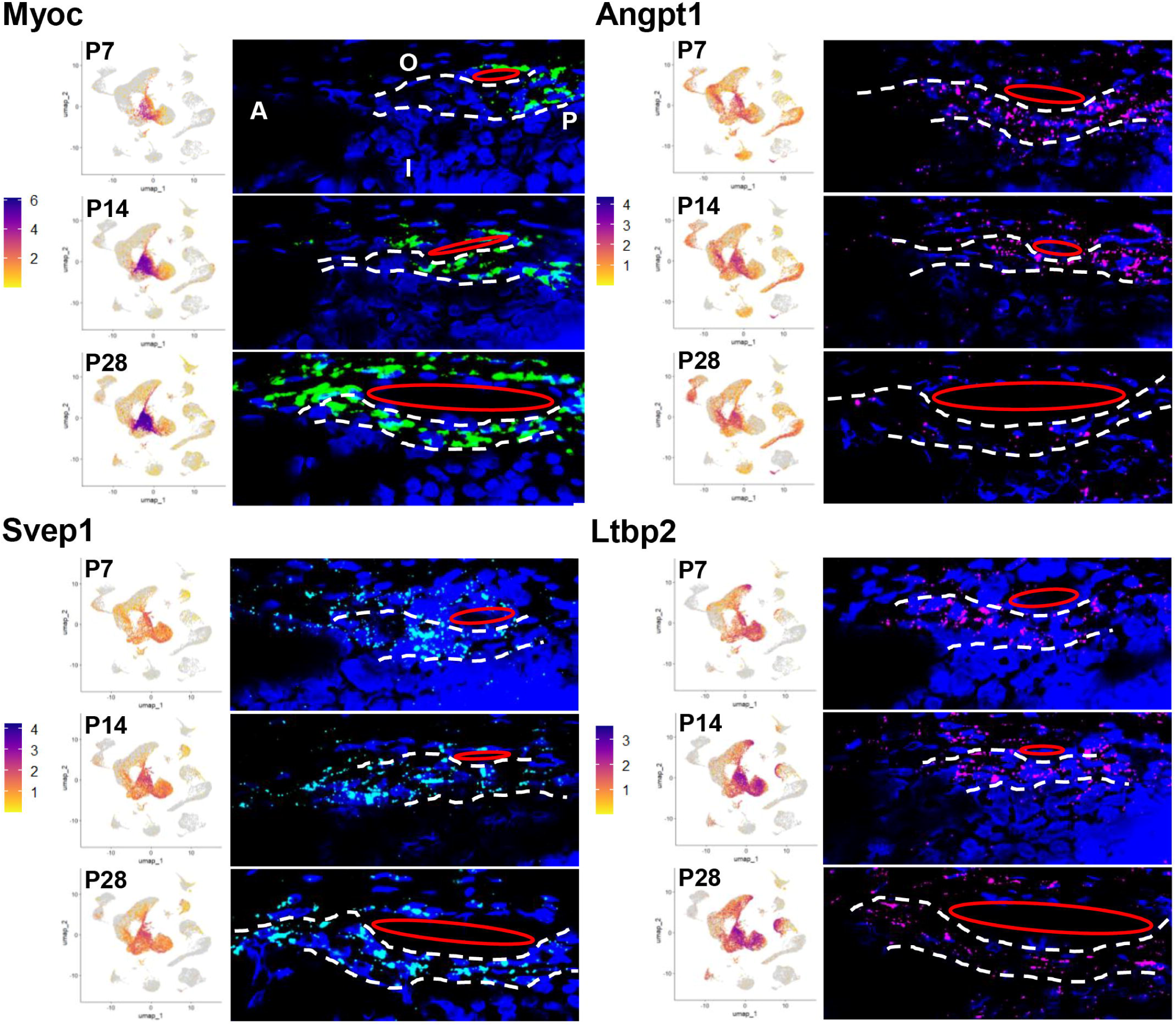
Limbal expression of *Myoc*, *Angpt1*, *Svep1*, and *Ltbp2* in TMCs (fibroblast-like cells, C1). Gene expression derived from scRNAseq data (normalized log2 fold change, left) alongside tissue sections stained by ISH (right) at each age. All ISH images are oriented the same as the first panel with the anterior side (A, closer to the pupil) on the left, posterior side (P, closer to the retina) on the right, the outer side of the eye (O) towards the top, and the inner side (I) towards the bottom. TM location is highlighted between dashed white lines and the position of SC is displayed as a red ellipse.

C1 cells were isolated and reclustered into four fibroblast-like (FBL0-FBL3) subtypes (Fig S7), and their identities inferred from established marker genes, as described in Supporting Information (“Identification of fibroblast-like subtypes” in S1 Appendix and Fig S7; heatmap of top differentially expressed genes between FBL clusters provided in Fig S8). Putative TMC-like cells were identified within FBL0, isolated, and the 10,037 cells re-clustered into seven groups (TMC0-TMC6, Fig 6; heatmap of top differentially expressed genes between TMC clusters provided in Fig S9). The TMC cluster data correlated with ISH studies in tissue sections, which showed expression of the pan-TM marker *Myoc* increased progressively with tissue maturation, starting posterior to the rudimentary SC and spreading throughout the TM as it developed (Fig S10). Conversely the *Cyp1b1* gene, in which mutations account for most cases of primary congenital glaucoma, was expressed more highly in P7 and P14 TMCs than at P31 (Fig S11).

**Fig 6.**
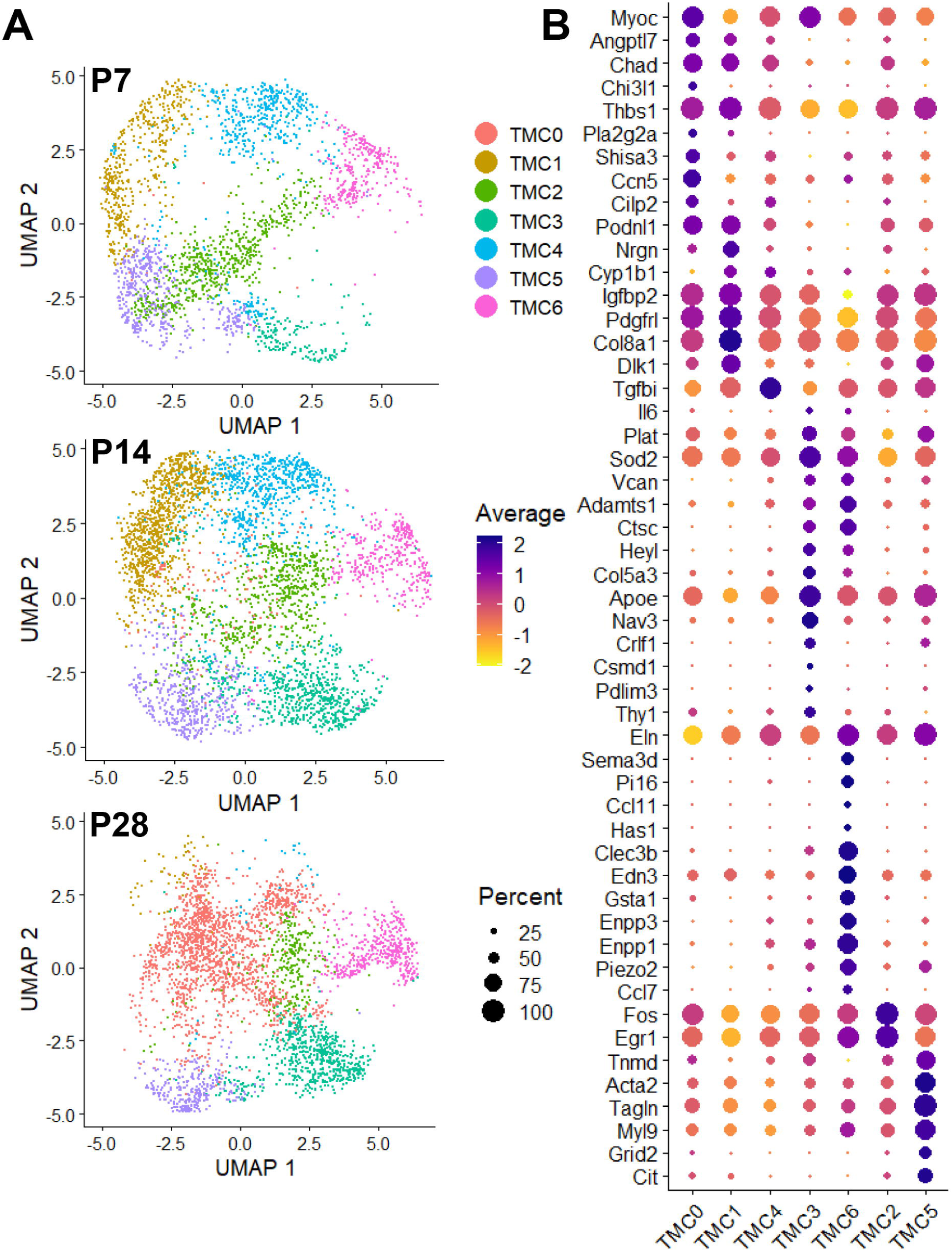
Cellular subtypes and associated gene expression within the developing trabecular meshwork cell population. (A) UMAP plot of seven TMC-like subtypes (TMC0-TMC6) derived from FBL0 at each age. (B) Dot plot of key marker gene expression distinguishing TMC-like subtypes.

To investigate the developmental trajectory of these subpopulations, we analyzed changes in cell abundance across developmental timepoints. Cell populations in TMC1, TMC2, and TMC4 decreased over time, suggesting they represent progenitor or transitional states (Fig 6). Notably, TMC2 cells exhibited a marked shift in UMAP position, indicating broad changes in expression profile as the TM developed. TMC5 cells largely maintained their numbers, while TMC6 cells gradually expanded - both accompanied by dynamic changes in gene expression. In contrast, TMC0 and TMC3 expanded significantly during mid-to-late developmental stages, which implicated them as terminally differentiated TMCs. Minimal expression of proliferation (*Mki67*, *Top2a*) and apoptosis (*Casp3*, *Casp7*^68^) markers suggested cell state differentiation as the major cause of the cluster changes.

The largest terminal cluster, TMC0, showed the strongest *Myoc* expression but contained few cells at P7 (Fig 6). This population expanded substantially over time and expressed hallmark markers of the JCT, including *Chad*, *Angptl7*, *Chi3l1*, and *Thbs1*.^66^ Additional genes enriched in TMC0 cells included *Pla2g2a*, *Shisa3*, *Ccn5*, *Cilp2*, *Podnl1*, and *Nrgn*. ISH studies of *Chad* detected strong expression in corneal and limbal stroma and confirmed weaker expression within a subset of the TM that localized more closely to the JCT at P31 (Fig S12).

The second and third largest terminal clusters, TMC3 and TMC6, expressed key markers of the uveal and corneoscleral TM layers, including *Plat* (tissue-type plasminogen activator), *Il6* (interleukin-6), and *Sod2* (superoxide dismutase 2).^60–62^ The clusters also expressed *Vcan* (versican), which binds HA to form large proteoglycan aggregates and plays a role in modulating AH outflow resistance in the JCT^69,70^, and the metalloprotease *Adamts1*, which cleaves versican into its bioactive form.^71^ These data suggest that, like HA, versican is secreted by uveal and corneoscleral TMCs and transported into the JCT by aqueous flow.^63^ TMC3 and TMC6 cells also expressed *Gsn*, *Ctsc*, *Heyl*, and the TM lamellae-associated type V collagen (*Col5a3*^72^). ISH studies of *Gsn* detected moderate expression widely throughout the P7 and P14 tissues which became strongly enriched in the TM layers by P31 (Fig S13). Specifically, TMC3 cells could be distinguished by selective expression of *Apoe*, *Nav3*, *Crlf1*, *Csmd1*, and *Pdlim3*. ISH studies of *Apoe* confirmed selective expression within a subset of the TM (Fig S14). Conversely, TMC6 cells uniquely expressed HA synthase (*Has1*),^63,64^ TM-associated mechanosensitive ion channel (*Piezo2*),^73^ *Clec3b*, *Sema3d*, *Pi16*, *Ccl11*, *Edn3*, *Gsta1*, *Enpp3*, *Enpp1*, and *Ccl7*. ISH studies for *Clec3b* confirmed expression localized to a specific subset of the TM at P14 and P31 (Fig S15).

Significant TMC2 and TMC5 cell populations were present at all three developmental ages with subtle changes in cluster locations noted, reflecting overall gene expression shifts. TMC2 cells shared a similar profile to TMC0, that included JCT-like *Chad* and *Angptl7* expression, but with higher levels of *Fos* and *Egr1*. ISH studies of *Fos* localized specific expression to the posterior region of the TM at P7 and P14 that was adjacent to SC by P31 (Fig S16). In contrast, TMC5 cells displayed a transcriptional profile that included strong tendon (*Tnmd*^74^) and smooth muscle (*Acta2*, *Tagln*, *Myl9*, *Eln*) markers, as well as unique expression of *Grid2* and *Cit*. ISH studies of *Tnmd* detected highly specific expression throughout the TM at P7 and P14 that extended to adjacent stromal fibroblasts by P31 (Fig S17).

Finally, the two early-stage TM populations (TMC1 and TMC4) were significantly reduced by P28, which was consistent with progenitor cells contributing to TM formation. Both clusters expressed high levels of *Chad* and *Angptl7*, supporting their identity as JCT progenitors. Other genes expressed by TMC0/JCT cells were also detected at higher levels in TMC1 cells, such as *Igfbp2*, *Pdgfrl*, *Col8a1*, and *Nrgn*. ISH studies of *Igfbp2* confirmed strong expression throughout the early TM at P7 that persisted at P31 (Fig S18). Early stage TMC1 cells also shared expression of *Dlk1* with the later TMC2 and TMC5 cell types. Conversely, TMC4 cells were marked by higher levels of *Tgfbi*, a protein important for cell-ECM interactions,^75^ which ISH studies confirmed peak expression at P14 in TMCs located between SC and the ciliary body (Fig S19).

### Selective gene expression in SECs and TMCs

In total, this study identified 655 SC- and 10,037 TM-related cells that could be further subdivided into 13 distinct populations according to their distinguishing gene expression profiles. To recognize molecular pathways important for the development and function of SC and the TM, differential expression analyses were performed to identify genes with enhanced expression in these important cell types versus all other limbal cells. For the four main SEC subtypes (SEC0, SEC1, SEC2, and SEC4), 244 genes with human orthologs were selectively expressed in one or more of the subclusters. Genes selectively expressed by the proliferating BEC population (SEC3) were not included as these cells could not be definitively related to SC versus other developing blood vessels. Only *Hoxd8*, a transcription factor important for the differentiation of BECs into lymphatic-like endothelial cells,^76^ was selectively expressed in the proliferating SEC population (SEC5) compared to SEC3 cells. Its inclusion brought the total number of selectively upregulated SEC genes to 245. In total for all seven TMC subtypes, 153 genes with human orthologs were selectively upregulated in one or more of the subclusters. Three genes, *Ephb1*, *Lbp*, and *Sema3d*, were identified in subtypes of SECs and TMCs. Combined, this study identified 395 genes selectively upregulated in cell types associated with these two critical developing outflow pathway tissues (Fig 7).

**Fig 7.**
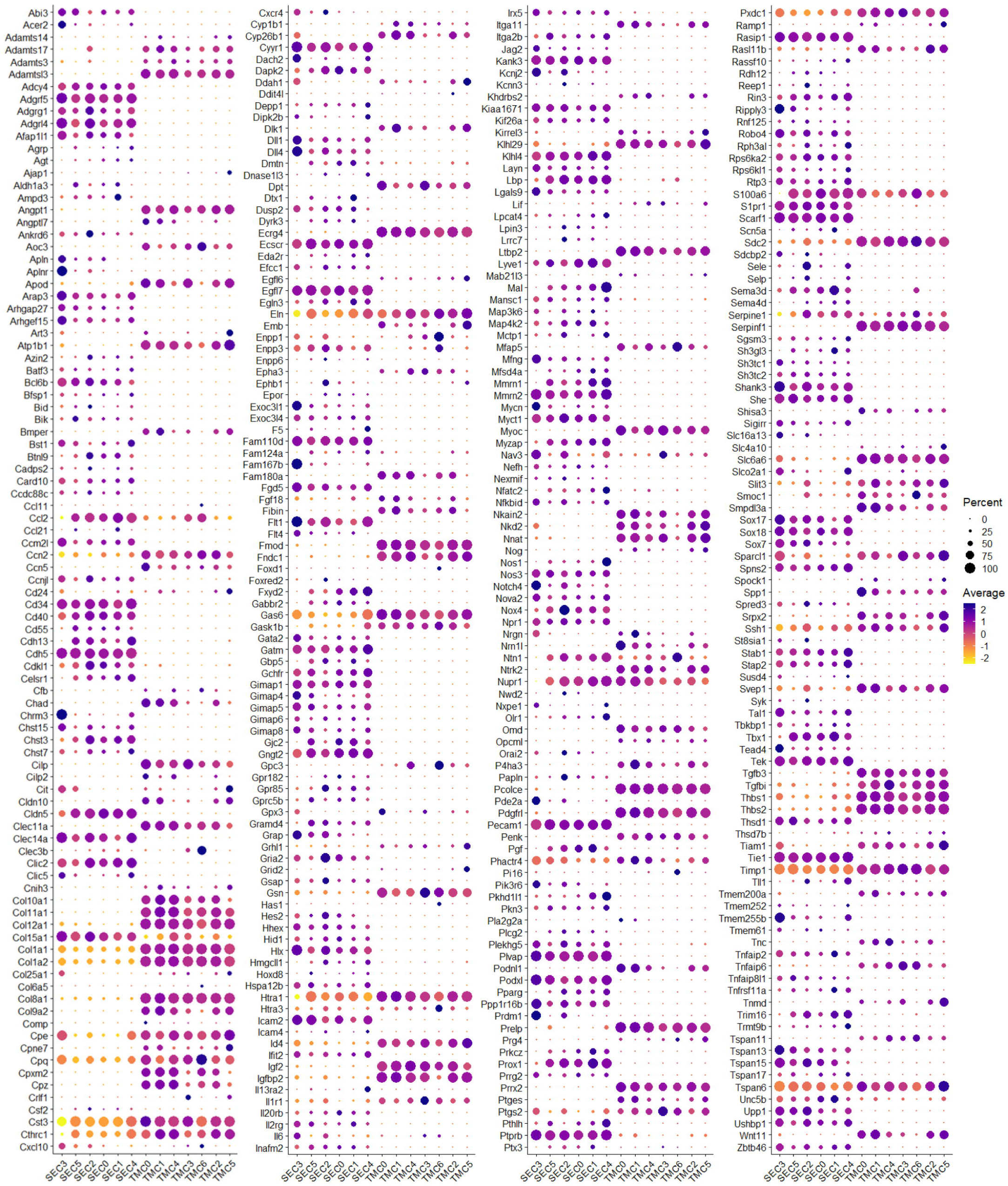
Dot plot visualization of the relative expression of 395 genes selectively expressed in SC- and TM-related cell clusters. Gene names are listed along the y-axis and cell type cluster names are shown along the x-axis; six SC- related clusters (SEC0-SEC5) and seven TM-related clusters (TMC0-TMC6). Average gene expression is relative to other SC and TM clusters.

### Gene expression associated with Mendelian disease

We sought to assess whether genes already associated with Mendelian forms of congenital glaucoma are expressed within our developing SC/TM dataset. Primary congenital glaucoma falls within a larger group of congenital glaucomas that includes syndromic forms encompassing non-ocular organ systems and developmental anomalies collectively referred to as anterior segment dysgenesis (ASD; OMIM 107250). ASD shows vast phenotypic variability of abnormalities affecting the iris, cornea, lens, TM, and SC,^77^ and are caused by mutations in many genes including *PAX6*, *FOXC1*, *PITX2*, *FOXE3*, *PITX3*, *PXDN*, *CPAMD8*, *B3GLCT*, and *COL4A1*.^77–79^ It can be difficult to diagnostically separate cases of primary congenital glaucoma from the other disorders when the clinical features outside of the drainage tissues are mild.^78,79^ Together, this study interrogated 44/45 genes previously associated with congenital glaucoma. One gene, *Guca1c*, could not be assessed in our data due to the lack of a rat ortholog. The data revealed 36/44 (82%) genes were expressed by at least one SC- or TM-related cell cluster (Fig 8).

**Fig 8.**
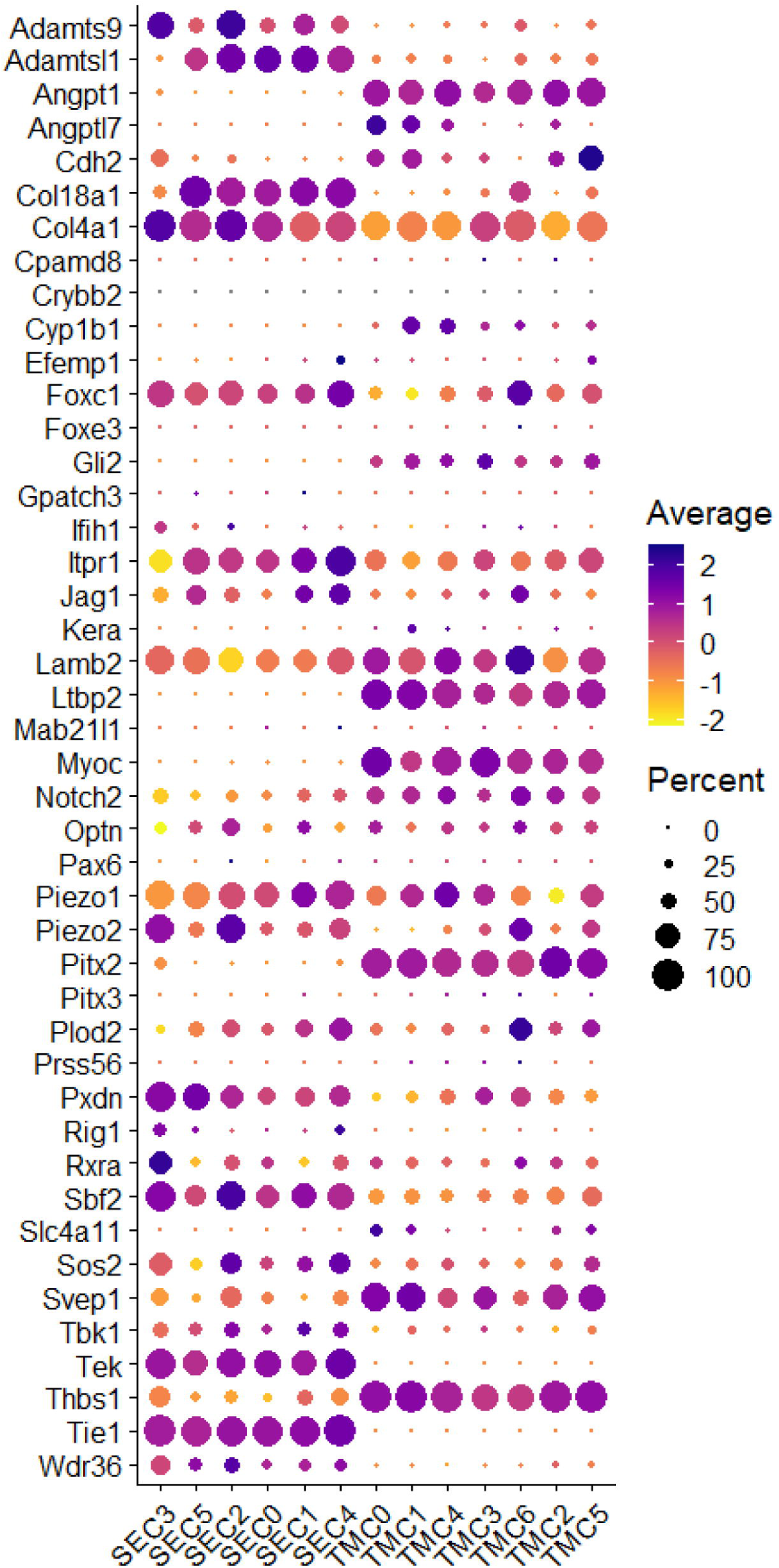
Dot plot visualization of the relative expression of 44 Mendelian congenital glaucoma-associated genes between SEC and TMC subtypes.

### Gene expression associated with complex disease

Genome-wide association studies (GWAS) have implicated hundreds of genes in the pathogenesis of glaucoma. Primary open-angle glaucoma (POAG), the leading cause of irreversible blindness worldwide, is characterized by late-onset optic nerve damage, progressive visual field loss, and typically elevated IOP despite open access of AH to the drainage tissues.^80,81^ In 2021, Gharahkhani et al. reported a multi-ethnic meta-analysis of GWAS data from 34,179 POAG cases and 349,321 controls, and identified 127 risk loci that collectively explained 9.4% of familial risk.^82^ Of these, 89/123 loci with available IOP data were significantly associated with IOP, which suggested shared genetic etiology. More recently, Han et al. (2023) reported the largest multitrait gene-based GWAS analysis (MTAG), which identified 355 genes significantly associated with POAG risk.^83^

Given the expected role of AHOP dysfunction in POAG, we examined the expression of these POAG- associated genes in our dataset. Genes were filtered to remove duplicates between the two GWAS lists, those without known protein-coding rat orthologs, or those lacking expression in our scRNAseq data. In cases where multiple protein-coding genes mapped to a single locus, all were included in the assessment. Together, 372 POAG-associated genes were interrogated across SC/TM-related cell clusters (Fig S20). Many of the genes demonstrated high expression in SECs and/or TMCs, but few were selectively expressed in just those cells versus other limbal cell types. Thirteen genes were selectively expressed at substantial levels by SECs, which were *Adam15*, *Cav1*, *Cav2*, *Clic5*, *Cttnbp2*, *Egln3*, *Emid1*, *Fam53b*, *Kctd20*, *Maff*, *Pthlh*, *Tes*, and *Thsd7a*. Fourteen genes were selectively expressed at robust levels by TMCs, which were *Angpt1*, *Col24a1*, *Efemp2*, *Itgb5*, *Ltbp1*, *Ltbp2*, *Ltbp3*, *Myoc*, *Nucb2*, *Pitx2*, *Rora*, *Svep1*, *Tcf7l2*, and *Tgfb3*. Two genes, *Pde7b* and *Plce1*, were selectively enriched at in populations of both SECs and TMCs.

## Discussion

This work has demonstrated the feasibility of the scRNAseq approach to broadly catalogue cell types and their associated gene expression profiles in the diminutive and complex tissues of the developing rat AHOP. This method has identified 655 VECs related to SC development, including 109 actively proliferating and 546 quiescent, as well as 10,037 cells related to the development of the TM. ISH studies reinforced the localization of these cell types to the TM/SC. Further refinement to specific regions, implicated by known marker gene expression, was restricted by the limited rat tissue size and low nuclear transcript signals obtained from thin ISH sections. As antibodies proven for rat protein detection become available, further whole tissue staining studies will enable a more detailed investigation.

Interrogation of 44 genes linked with Mendelian forms of congenital glaucoma revealed 36/44 (82%) were expressed by at least one SC- or TM-related cell cluster, validating the dataset’s relevance. The classification of cell types presented here is consistent with published studies such as the expression of *Angpt1*, *Angptl7*, *Cyp1b1*, *Ltbp2*, *Myoc*, *Piezo1*, *Pitx2*, *Svep1*, and *Thbs1* in the TM,^3,5–8,57,58,66,84,85^ and *Col4a1*, *Piezo2*, *Tek*, and *Tie1* in SC.^2,12,73,86^ Furthermore, for the eight genes which lacked TM/SC cluster expression, published findings describe expression and disease mechanisms that involve tissues outside of the AHOP or function during earlier developmental processes. Specifically, *CPAMD8* variants cause congenital glaucoma with anterior segment dysgenesis, myopia, and ectopia lentis, likely due to abnormal iris or lens morphology rather than SC/TM defects.^79,87^ *CRYBB2*, associated with congenital cataracts and glaucoma, is expressed in lens epithelial cells and retinal neurons, and directly prevents RGC loss.^88–90^ *FOXE3* and *PITX3*, both involved in early lens development, are linked to anterior segment dysgenesis, with lens-cornea or iris-lens adhesions obstructing aqueous outflow.^91–93^ *PRSS56* mutations cause posterior microphthalmia, leading to angle-closure glaucoma due to a shallow anterior chamber.^94^ *MAB21L1* mutations, associated with syndromic and ocular abnormalities including buphthalmos, impair anterior segment architecture in zebrafish, reducing outflow via fibrous masses between the lens and cornea.^95–98^ *PAX6* is essential for early differentiation of periocular mesenchyme, and its loss in mice leads to an undeveloped TM and absent SC – events occurring too early in development (P0-P4) to be captured in this study.^99–101^ Similarly, *GPATCH3* is implicated in congenital glaucoma, but its expression has only been observed in early zebrafish periocular mesenchyme and corneal endothelium.^102^

Unexpectedly, our studies revealed expression of *Lyve1* in rat SECs. This is in contrast to expression reported for SC in mice, where *Lyve1* expression is only associated with resident macrophages and not SECs.^12^ *Lyve1* expression is controlled by specific signaling pathways, such as those involving *Prox1*, and encodes a hyaluronan receptor that primarily helps regulate the transport of immune cells and fluid through lymphatic vessels.^103^ This function is distinct from that of blood vessels, which largely manage nutrient movement and gas exchange; consequently *Lyve1* is not expressed by BECs. However, SC is known to show hybrid characteristics of lymphatic and blood vasculature, expressing BEC-like *Cd34*, *Flt1*, *Nos3*, and LEC-like *Prox1*. Like lymphatic vessels, SC is highly involved in fluid homeostasis and immune cell trafficking from the eye, for which *Lyve1* may play a role. Our data is consistent with this function for *Lyve1* in the SC of rats with expression that is negligible in early SECs but upregulated as the cells mature and acquire a lymphatic-like state.

The identification of novel single gene causes of congenital glaucoma has not been forthcoming despite large-scale sequencing efforts. It is likely that there are many undiscovered genes in which strong-effect variants account for a limited proportion of the disease, but these are difficult to identify in individuals or small-sized patient cohorts. Utilizing the transcriptomic resource presented here, interrogation of variants in developing AHOP-expressed genes can now be prioritized. Importantly, 395 genes were detected as selectively upregulated during the development of the TM/SC. Noteworthy genes significantly upregulated by SC-related cells were *Agt*, *Ccl2*, *Cdkl1*, *Chst3*, *Clic2*, *Dapk2*, *Lbp*, *Mmrn1*, *Myzap*, *Tbx1*, and *Trim16*. For TM-related cell types, prominent genes included *Adamtsl3*, *Epha3*, *Fmod*, *Itga11*, *Nkd2*, *Phactr4*, and *Prelp*. These genes are likely critical for the development and/or function of the AHOP and warrant further investigation. We propose prioritization of these genes during patient genome analyses as they represent strong novel candidates for congenital glaucoma-causing variants.

It is probable that a proportion of the unresolved disease may not follow single gene/Mendelian inheritance but instead involve complex polygenic mechanisms. Hundreds of genes have already been associated with glaucoma endophenotypes by GWAS, and investigators are now utilizing this knowledge to generate polygenic risk scores (PRS) – an approach that estimates an individual’s disease risk based on their overall genetic makeup. This transcriptomic dataset will help adjust the weighting of variants in genes more highly expressed during AHOP tissue development, which in turn will enable the generation of PRS estimates tailored towards early-onset forms of glaucoma.

## Materials and methods

### Ocular tissue acquisition

All animal procedures performed in the study were in accordance with the ARVO Statement for Use of Animals in Ophthalmic Vision and Research. The ethical principles established by the National Institutes of Health Guide for the Care and Use of Laboratory Animals (revised 2011) were followed. The research protocol was ethically reviewed and approved by the University of Wisconsin–Madison School of Medicine and Public Health Institutional Animal Care and Use Committee.

Fischer 344 (F344/NCrl, Charles River Laboratories) rats, an albino, inbred strain of the common rat (*Rattus norvegicus*), were bred using standard husbandry practices. Staggered breeding pairs were established to collect all 18 limbal tissue strips within a three-day window. To minimize acquisition bias, dissections were performed in six groups, systematically changing the ascertainment order across genders and developmental age (S2 Table). Eyes were removed and immediately placed in ice-cold Hanks’ Balanced Salt Solution (HBSS) without calcium. Extraocular muscle was removed, and a scalpel was used to make an incision in the sclera just posterior to the limbus. Fine spring scissors were used to cut circumferentially around the globe to isolate the anterior cup. The lens and retina were removed, and the anterior cup was bisected through the pupil with a scalpel. The iris was trimmed to retain only a small region adjacent to the ciliary body. The opaque sclera was dissected away just posterior to the ciliary body, and the clear cornea was excised immediately anterior to the limbus. The resulting limbal tissue strips were maintained in ice-cold calcium-free HBSS until all three samples in the group were ready for simultaneous dissociation.

Limbal tissue strips were enzymatically dissociated into single cells using a two-step protocol. First, the tissue was digested with Collagenase Type 1 (CLSS-1, 10 mg/mL; Worthington) in HBSS supplemented with calcium and magnesium at 37°C for 1.5 hours, with gentle inversion every 15 minutes. Following centrifugation at 300 RCF for 5 minutes (a condition used for all subsequent centrifugation steps), the pellet was further digested with TrypLE at 37°C for 20 minutes. During this step, samples were inverted at 5 and 10 minutes and gently pipetted with a wide-bore P1000 tip at 15 and 20 minutes to enhance dissociation.

After centrifugation, the largely dissociated cells were resuspended in ice-cold DMEM supplemented with 10% FBS. Residual tissue was gently dissociated by pipetting with a regular-bore P1000 tip. The cell suspension was filtered through a 70 μm Flowmi tip, centrifuged, and resuspended in 100 μL by pipetting with a wide-bore P200 tip, followed by a regular-bore P200 tip. The sample was then passed through a 40 μm Flowmi tip. Cell count and viability were assessed with a LUNA FX7, ensuring 11,200 viable single cells (>75% viability) were prepared for each sample. Cells were loaded into a well of a Chromium Next GEM Chip G, and Single-Cell 3ʹ Gene Expression libraries generated following the manufacturer’s protocol (Chromium Next GEM Single-Cell 3ʹ Reagent Kit v3.1 User Guide, Rev D).

### Single-cell RNA sequencing and bioinformatic processing

Eighteen dissociated tissue samples were independently processed through 18 GEM wells for 3’ cDNA library construction. A preliminary low-read-deep sequencing run was conducted on an Illumina MiSeq to estimate the total cell counts and reads per cell across libraries. Quality control was performed using UMI-tools.^104^ The 18 libraries were balanced, pooled, and deeply sequenced on a single S4 flow cell using the Illumina NovaSeq platform. Demultiplexing was performed with Cell Ranger’s mkfastq command, utilizing Illumina’s bcl2fastq wrapper. Subsequent alignment, filtering, and UMI counting for each sample was completed using the count command, in accordance with 10x Genomics’ documentation. Reads were aligned to the latest rat genome reference (GRCr8, GenBank GCA_036323735.1, Jan 31, 2024) with the mitochondrial reference contig added from an earlier genome build (Rnor_6.0). Alignments included introns to enhance expression sensitivity and excluded pseudogenes to improve mapping quality.

Cell Ranger generated barcode matrices for each sample, which were aggregated using the aggr command. Unique GEM well suffixes were appended to barcodes to retain sample identity. Aggregation included depth normalization, ensuring equal average read depth per cell across samples to avoid sequencing depth-related artifacts. As all libraries were sequenced together in a single flow cell, no batch correction was necessary. The final aggregated matrix comprised raw normalized expression data from 126,787 gel bead droplets that were suitable for comparative analyses.

### Bioinformatic analyses

Data analysis was conducted primarily with the Seurat v 5.0.1 R package.^105^ A Seurat object was created from the aggregated barcode matrix, including genes expressed in at least five cells. Low-quality cells were filtered out based on the following criteria: UMI count < 3000, fewer than 1200 genes expressed, or mitochondrial gene content > 10%. Mitochondrial content was calculated across 13 protein-coding mitochondrial genes (Mt-co1, Mt-nd6, Mt-atp6, Mt-co2, Mt-co3, Mt-nd3, Mt-nd1, Mt-nd4, Mt-nd5, Mt- atp8, Mt-nd2, Mt-nd4l, Mt-cyb). This filtering reduced the initial 126,787 cells to 90,274 high-quality cells.

Multiplets were detected and removed (∼4%) using scDblFinder, yielding a final dataset of 86,653 single cells and 29,626 genes. Expression data were normalized using Seurat’s NormalizeData function, which scales gene counts per cell to 10,000, followed by log transformation.

For dimensional reduction and clustering, Seurat’s SCTransform function was used to normalize, scale, and identify variable genes. Principal component analysis was performed using RunPCA, and an elbow plot determined that the first 18 principal components captured most variance. A Shared Nearest Neighbor (SNN) graph was constructed with FindNeighbors, and clustering was done using FindClusters (resolution = 0.1). A two-dimensional Uniform Manifold Approximation and Projection (UMAP) was generated with RunUMAP, visualizing 13 distinct cell-type clusters according to each cell’s overall gene expression similarity.

Differential gene expression was assessed using the FindMarkers command in conjunction with the Model-based Analysis of Single-cell Transcriptomics (MAST) algorithm - a statistical tool that models gene expression via a zero-inflated hurdle model to account for sparsity specifically in single-cell datasets.^106^ For TMCs, the criteria for selective expression were a greater than two log2 fold change upregulation in more than 20% of cells within the TMC subcluster versus all other limbal cells and robust expression (significant UMI counts or highly cluster-specific UMIs at low level). For SECs, as all the VECs are very different to other limbal cells, a more stringent criteria needed to be implemented with expression thresholds set at a greater than two log2 fold change upregulation in more than 30% of cells within the SEC subcluster versus all other limbal cells, less than 10% expression in other limbal cells, and robust expression (significant UMI counts or highly cluster-specific UMIs at low level).

UMAP and HeatMap plots were generated using the standard Seurat package commands DimPlot and DoHeatMap, respectively. Customized DotPlot and FeaturePlot (individual gene expression overlayed on UMAP) images were generated using the DotPlot_scCustom and FeaturePlot_scCustom commands within the scCustomize package (https://doi.org/10.5281/zenodo.5706430).^107^

### Whole mount fluorescent immunohistochemistry

Fischer rats were euthanized via decapitation at P4 and P7 or CO2 gas inhalation at P14 and P28. Eyes were rapidly enucleated and placed in ice-cold 1x phosphate buffered saline pH 7.4 (PBS). Eyes were punctured with a 30G needle posterior to limbus, and immersion fixed in 4% paraformaldehyde (PFA) in PBS for 10 minutes at 4^°^C. Following fixation, tissues were washed in 70% ice-cold molecular biology grade ethanol (EtOH) and stored in fresh 70% EtOH at 4^°^C. For staining, eyes were transferred to a Kimwipe soaked in ice-cold PBS for dissection. Using spring scissors and fine tip forceps, extraneous tissues such as the conjunctiva, muscle, and tendon were removed, eyes bisected, and lenses extracted. Anterior eye cups were transferred into a dish containing ice-cold PBS where the retina and ciliary body were removed. The tissues were placed into blocking/staining buffer (5% normal donkey serum (Cat. 017-000-121, Jackson ImmunoResearch Labs) 0.5% triton x-100, and 0.01% sodium azide in PBS) on a nutator overnight at 4^°^C. Blocking buffer was replaced with primary antibodies diluted in staining buffer and incubated on a nutator at 4^°^C for approximately 24 hours. Mouse anti-CD31/PECAM1 (Cat. MA1- 80069, Invitrogen) was diluted 1:100, rabbit anti-LYVE-1 (Cat. 67538, Cell Signaling) at 1:500, and goat anti-AIF-1 (Cat. NB100-1028, Novus) at 1:100. Primary antibodies were removed, and eye cups washed three times in PBS containing 0.01% Tween 20 for one hour at room temperature (RT).

Secondary antibodies diluted in staining buffer were added and tissues incubated on a nutator at 4^°^C for approximately 24 hours. Donkey anti-mouse Alexa 405 (Cat. A48257, Invitrogen) was diluted at 1:500, donkey anti-rabbit Alexa 546 (Cat. A10040, Invitrogen) at 1:500, and donkey anti-goat Alexa 647 (Cat. 705-605-147, Jackson ImmunoResearch Labs) at 1:500. Following secondary antibody removal, tissues were washed four times with PBS containing 0.01% Tween 20 for 1 hour at RT. Eye cups were transferred into cold PBS and a series of relief cuts were made towards the center that resulted in a flower petal appearance. Tissues were placed on glass microscope slides with the cornea side down, excess fluid removed, and mounted (Shandon-Mount, Cat. 1900331, Epredia) with #1.5 coverslips. Slides were dried for >16 hours at 4^°^C and imaged with a Nikon A1R confocal microscope using a 20x objective and 23 um pinhole (1 Aery unit). A 75 um Z-stack of 1 um slices was collected, which encompassed all of SC.

### *In situ* hybridization (RNAscope)

Rats were euthanized by decapitation (P7) or CO2 gas inhalation (P14 and P31). The eyes were rapidly enucleated and transferred into ice-cold 4% PFA in 1x PBS, pH 7.4 (4% PFA). Using a scalpel, a large incision was made posteriorly to the limbus to allow rapid PFA access to internal eye structures. Tissues were placed into fresh ice-cold 4% PFA at 4°C for 2 hours. Eyes were dissected in 4% PFA on ice, removing extraneous muscle and tendon, bisecting posterior to the limbus, and removing the lens. Fixation of the anterior eye cups continued in 4% PFA at 4°C for a total of 20 hours. Tissues were processed through an alcohol series, xylene, and embedded in paraffin. Blocks were stored at 4°C in a sealed container with desiccant. Paraffin blocks were sectioned at 5 µm thickness and allowed to dry overnight at RT. Sections were processed for RNAscope following the manufacturers protocol (User Manual for RNAscope Multiplex Fluorescent Reagent Kit v2, UM 323100/rev B/Effective Date: 10/11/2022). Mild conditions were used for antigen retrieval (10 minutes) and standard conditions for the protease treatment (30 minutes). Hybridization was performed in a HybEZ II oven (Cat. 321710/321720). Details of RNAscope probes are provided in S3 Table, which were used in conjunction with the following Opal dye fluorophores (Akoya Biosciences): Opal 520 (Cat. FP1487001KT), Opal 570 (Cat. FP1488001KT), Opal 620 (Cat. FP1495001KT), and Opal 650 (Cat. FP1496001KT). Images were collected with a Nuance Multispectral Imaging System including an Olympus BX43 microscope with 40x objective and Perkin Elmer Nuance software v.3.0.2).

### Data availability

The raw data files from this study have been deposited in the Sequence Read Archive of the National Center for Biotechnology Information (accession no. PRJNA1311507).

## Acknowledgments

This study was funded by the National Eye Institute (NEI) R21 EY034251 (to ST), and supported by the Margaret Emma Williams Trust Fund to the University of Wisconsin-Madison Department of Ophthalmology and Visual Sciences, an Unrestricted Grant from Research to Prevent Blindness, Inc. to the University of Wisconsin-Madison Department of Ophthalmology and Visual Sciences, and a Core Grant for Vision Research from the National Institutes for Health (NIH) to the University of Wisconsin- Madison (P30 EY016665).

The authors utilized the University of Wisconsin-Madison Biotechnology Center’s Gene Expression Center (Research Resource Identifier – RRID:SCR_017757) for 10X Genomics scRNAseq library preparation, DNA Sequencing Facility (RRID:SCR_017759) for Illumina sequencing, and Bioinformatics Core Facility (Research Resource Identifier – RRID:SCR_017799) for initial processing of raw scRNAseq data. The authors wish to thank the University of Wisconsin Translational Research Initiatives in Pathology (TRIP) Laboratory, supported by the UW Department of Pathology and Laboratory Medicine, UWCCC (P30 CA014520) and the Office of The Director- NIH (S10 OD023526) for its histology and RNAscope services, and for the use of their facilities. Confocal imaging was facilitated by the University of Wisconsin Optical Imaging Core, and we thank Lance A Rodenkirch for microscopy support.

We thank the department chair, Dr. Terri L. Young, for provision of technical support staff, laboratory space, and start-up funding to Dr. Tompson. The authors would also like to thank Dr. Abigail Radcliff in the laboratory of Dr. Jyoti Watters (UW School of Veterinary Medicine), and Miranda Pamperin, Riley Brutto, Devin Larson, and John Michael in the laboratory of Dr. Edwin Chapman (UW Neuroscience) for providing rat tissues for preliminary studies. We also thank Dr. Stephanie L. Nay for critical review of the manuscript.

## References

1. García-Antón MT, Salazar JJ, de Hoz R, et al. Goniodysgenesis variability and activity of CYP1B1 genotypes in primary congenital glaucoma. PLoS One. 2017;12(4):e0176386. doi:10.1371/journal.pone.0176386

2. Souma T, Tompson SW, Thomson BR, et al. Angiopoietin receptor TEK mutations underlie primary congenital glaucoma with variable expressivity. J Clin Invest. Jul 1 2016;126(7):2575–2587. doi:10.1172/jci85830

3. Thomson BR, Souma T, Tompson SW, et al. Angiopoietin-1 is required for Schlemm’s canal development in mice and humans. J Clin Invest. Dec 1 2017;127(12):4421–4436. doi:10.1172/jci95545

4. Kuehn MH, Lipsett KA, Menotti-Raymond M, et al. A Mutation in LTBP2 Causes Congenital Glaucoma in Domestic Cats (Felis catus). PLoS One. 2016;11(5):e0154412. doi:10.1371/journal.pone.0154412

5. van Zyl T, Yan W, McAdams A, et al. Cell atlas of aqueous humor outflow pathways in eyes of humans and four model species provides insight into glaucoma pathogenesis. Proc Natl Acad Sci U S A. May 12 2020;117(19):10339–10349. doi:10.1073/pnas.2001250117

6. van Zyl T, Yan W, McAdams AM, Monavarfeshani A, Hageman GS, Sanes JR. Cell atlas of the human ocular anterior segment: Tissue-specific and shared cell types. Proc Natl Acad Sci U S A. Jul 19 2022;119(29):e2200914119. doi:10.1073/pnas.2200914119

7. Patel G, Fury W, Yang H, et al. Molecular taxonomy of human ocular outflow tissues defined by single-cell transcriptomics. Proc Natl Acad Sci U S A. Jun 9 2020;117(23):12856–12867. doi:10.1073/pnas.2001896117

8. Thomson BR, Liu P, Onay T, et al. Cellular crosstalk regulates the aqueous humor outflow pathway and provides new targets for glaucoma therapies. Nat Commun. Oct 18 2021;12(1):6072. doi:10.1038/s41467-021-26346-0

9. Balasubramanian R, Kizhatil K, Li T, et al. Transcriptomic profiling of Schlemm’s canal cells reveals a lymphatic-biased identity and three major cell states. Elife. Oct 18 2024;13 doi:10.7554/eLife.96459

10. Stamer WD, Clark AF. The many faces of the trabecular meshwork cell. Exp Eye Res. May 2017;158:112–123. doi:10.1016/j.exer.2016.07.009

11. Fuchshofer R, Welge-Lüssen U, Lütjen-Drecoll E, Birke M. Biochemical and morphological analysis of basement membrane component expression in corneoscleral and cribriform human trabecular meshwork cells. Invest Ophthalmol Vis Sci. Mar 2006;47(3):794–801. doi:10.1167/iovs.05-0292

12. Kizhatil K, Ryan M, Marchant JK, Henrich S, John SW. Schlemm’s canal is a unique vessel with a combination of blood vascular and lymphatic phenotypes that forms by a novel developmental process. PLoS Biol. Jul 2014;12(7):e1001912. doi:10.1371/journal.pbio.1001912

13. Park DY, Lee J, Park I, et al. Lymphatic regulator PROX1 determines Schlemm’s canal integrity and identity. J Clin Invest. Sep 2014;124(9):3960–3974. doi:10.1172/jci75392

14. Braakman ST, Moore JE, Jr., Ethier CR, Overby DR. Transport across Schlemm’s canal endothelium and the blood-aqueous barrier. Exp Eye Res. May 2016;146:17–21. doi:10.1016/j.exer.2015.11.026

15. Remé C, Urner U, Aeberhard B. The development of the chamber angle in the rat eye. Morphological characteristics of developmental stages. Graefes Arch Clin Exp Ophthalmol. 1983;220(3):139–153. doi:10.1007/bf02175946

16. Aspelund A, Tammela T, Antila S, et al. The Schlemm’s canal is a VEGF-C/VEGFR-3- responsive lymphatic-like vessel. J Clin Invest. Sep 2014;124(9):3975–3986. doi:10.1172/jci75395

17. Lang AJ, Mirski SE, Cummings HJ, Yu Q, Gerlach JH, Cole SP. Structural organization of the human TOP2A and TOP2B genes. Gene. Oct 23 1998;221(2):255–266. doi:10.1016/s0378-1119(98)00468-5

18. Gerdes J, Lemke H, Baisch H, Wacker HH, Schwab U, Stein H. Cell cycle analysis of a cell proliferation-associated human nuclear antigen defined by the monoclonal antibody Ki-67. J Immunol. Oct 1984;133(4):1710–1715.

19. Chakarov S, Lim HY, Tan L, et al. Two distinct interstitial macrophage populations coexist across tissues in specific subtissular niches. Science. Mar 15 2019;363(6432)doi:10.1126/science.aau0964

20. Köhler C. Allograft inflammatory factor-1/Ionized calcium-binding adapter molecule 1 is specifically expressed by most subpopulations of macrophages and spermatids in testis. Cell Tissue Res. Nov 2007;330(2):291–302. doi:10.1007/s00441-007-0474-7

21. Hsieh M, Huang PJ, Chou PY, et al. Carbonic Anhydrase VIII (CAVIII) Gene Mediated Colorectal Cancer Growth and Angiogenesis through Mediated miRNA 16-5p. Biomedicines. Apr 29 2022;10(5)doi:10.3390/biomedicines10051030

22. Yukiura H, Kano K, Kise R, Inoue A, Aoki J. LPP3 localizes LPA6 signalling to non-contact sites in endothelial cells. J Cell Sci. Nov 1 2015;128(21):3871–3877. doi:10.1242/jcs.172098

23. Icking A, Matt S, Opitz N, Wiesenthal A, Müller-Esterl W, Schilling K. NOSTRIN functions as a homotrimeric adaptor protein facilitating internalization of eNOS. J Cell Sci. Nov 1 2005;118(Pt 21):5059–5069. doi:10.1242/jcs.02620

24. Norrmén C, Ivanov KI, Cheng J, et al. FOXC2 controls formation and maturation of lymphatic collecting vessels through cooperation with NFATc1. J Cell Biol. May 4 2009;185(3):439–457. doi:10.1083/jcb.200901104

25. Valdembri D, Regano D, Maione F, Giraudo E, Serini G. Class 3 semaphorins in cardiovascular development. Cell Adh Migr. Nov 2016;10(6):641–651. doi:10.1080/19336918.2016.1212805

26. Aghajanian H, Choi C, Ho VC, Gupta M, Singh MK, Epstein JA. Semaphorin 3d and semaphorin 3e direct endothelial motility through distinct molecular signaling pathways. J Biol Chem. Jun 27 2014;289(26):17971–17979. doi:10.1074/jbc.M113.544833

27. Lee DJ, Kim P, Kim HY, et al. MAST4 regulates stem cell maintenance with DLX3 for epithelial development and amelogenesis. Exp Mol Med. Jul 2024;56(7):1606–1619. doi:10.1038/s12276-024-01264-5

28. Matsui J, Wakabayashi T, Asada M, Yoshimatsu K, Okada M. Stem cell factor/c-kit signaling promotes the survival, migration, and capillary tube formation of human umbilical vein endothelial cells. J Biol Chem. Apr 30 2004;279(18):18600–18607. doi:10.1074/jbc.M311643200

29. Koch AE, Halloran MM, Haskell CJ, Shah MR, Polverini PJ. Angiogenesis mediated by soluble forms of E-selectin and vascular cell adhesion molecule-1. Nature. Aug 10 1995;376(6540):517–519. doi:10.1038/376517a0

30. Buschman MD, Bromann PA, Cejudo-Martin P, Wen F, Pass I, Courtneidge SA. The novel adaptor protein Tks4 (SH3PXD2B) is required for functional podosome formation. Mol Biol Cell. Mar 2009;20(5):1302–1311. doi:10.1091/mbc.e08-09-0949

31. Hubmacher D, Schneider M, Berardinelli SJ, et al. Unusual life cycle and impact on microfibril assembly of ADAMTS17, a secreted metalloprotease mutated in genetic eye disease. Sci Rep. Feb 8 2017;7:41871. doi:10.1038/srep41871

32. Craige SM, Chen K, Pei Y, et al. NADPH oxidase 4 promotes endothelial angiogenesis through endothelial nitric oxide synthase activation. Circulation. Aug 9 2011;124(6):731–740. doi:10.1161/circulationaha.111.030775

33. Watanabe K, Hasegawa Y, Yamashita H, et al. Vasohibin as an endothelium-derived negative feedback regulator of angiogenesis. J Clin Invest. Oct 2004;114(7):898–907. doi:10.1172/jci21152

34. Wick N, Saharinen P, Saharinen J, et al. Transcriptomal comparison of human dermal lymphatic endothelial cells ex vivo and in vitro. Physiol Genomics. Jan 17 2007;28(2):179–192. doi:10.1152/physiolgenomics.00037.2006

35. Rousseau S, Houle F, Landry J, Huot J. p38 MAP kinase activation by vascular endothelial growth factor mediates actin reorganization and cell migration in human endothelial cells. Oncogene. Oct 1997;15(18):2169–2177. doi:10.1038/sj.onc.1201380

36. Zhu Q, Kim YH, Wang D, Oh SP, Luo K. SnoN facilitates ALK1-Smad1/5 signaling during embryonic angiogenesis. J Cell Biol. Sep 16 2013;202(6):937–950. doi:10.1083/jcb.201208113

37. Mannan A, Dhiamn S, Garg N, Singh TG. Pharmacological modulation of Sonic Hedgehog signaling pathways in Angiogenesis: A mechanistic perspective. Dev Biol. Dec 2023;504:58–74. doi:10.1016/j.ydbio.2023.09.009

38. Elbitar S, Renard M, Arnaud P, et al. Pathogenic variants in THSD4, encoding the ADAMTS-like 6 protein, predispose to inherited thoracic aortic aneurysm. Genet Med. Jan 2021;23(1):111–122. doi:10.1038/s41436-020-00947-4

39. Gunn MD, Tangemann K, Tam C, Cyster JG, Rosen SD, Williams LT. A chemokine expressed in lymphoid high endothelial venules promotes the adhesion and chemotaxis of naive T lymphocytes. Proc Natl Acad Sci U S A. Jan 6 1998;95(1):258–263. doi:10.1073/pnas.95.1.258

40. Vaahtomeri K, Moussion C, Hauschild R, Sixt M. Shape and Function of Interstitial Chemokine CCL21 Gradients Are Independent of Heparan Sulfates Produced by Lymphatic Endothelium. Front Immunol. 2021;12:630002. doi:10.3389/fimmu.2021.630002

41. Choi D, Park E, Jung E, et al. Laminar flow downregulates Notch activity to promote lymphatic sprouting. J Clin Invest. Apr 3 2017;127(4):1225–1240. doi:10.1172/jci87442

42. Choi D, Park E, Yu RP, et al. Piezo1-Regulated Mechanotransduction Controls Flow-Activated Lymphatic Expansion. Circ Res. Jul 8 2022;131(2):e2–e21. doi:10.1161/circresaha.121.320565

43. Zhao Y, Lin S. Essential role of SH3-domain GRB2-like 3 for vascular lumen maintenance in zebrafish. Arterioscler Thromb Vasc Biol. Jun 2013;33(6):1280–1286. doi:10.1161/atvbaha.112.301025

44. Black JA, Waxman SG. Noncanonical roles of voltage-gated sodium channels. Neuron. Oct 16 2013;80(2):280–291. doi:10.1016/j.neuron.2013.09.012

45. Wasik A, Ratajczak-Wielgomas K, Badzinski A, Dziegiel P, Podhorska-Okolow M. The Role of Periostin in Angiogenesis and Lymphangiogenesis in Tumors. Cancers (Basel). Aug 30 2022;14(17)doi:10.3390/cancers14174225

46. De Luca F, Kha M, Swärd K, Johansson ME. Identification of ARMH4 and WIPF3 as human podocyte proteins with potential roles in immunomodulation and cytoskeletal dynamics. PLoS One. 2023;18(1):e0280270. doi:10.1371/journal.pone.0280270

47. Vreeken D, Bruikman CS, Stam W, et al. Downregulation of Endothelial Plexin A4 Under Inflammatory Conditions Impairs Vascular Integrity. Front Cardiovasc Med. 2021;8:633609. doi:10.3389/fcvm.2021.633609

48. Morishita T, Tsutsui M, Shimokawa H, et al. Vasculoprotective roles of neuronal nitric oxide synthase. Faseb j. Dec 2002;16(14):1994–1996. doi:10.1096/fj.02-0155fje

49. Kuek V, Yang Z, Chim SM, et al. NPNT is Expressed by Osteoblasts and Mediates Angiogenesis via the Activation of Extracellular Signal-regulated Kinase. Sci Rep. Oct 26 2016;6:36210. doi:10.1038/srep36210

50. Simonson B, Chaffin M, Hill MC, et al. Single-nucleus RNA sequencing in ischemic cardiomyopathy reveals common transcriptional profile underlying end-stage heart failure. Cell Rep. Feb 28 2023;42(2):112086. doi:10.1016/j.celrep.2023.112086

51. Arystarkhova E. Beneficial Renal and Pancreatic Phenotypes in a Mouse Deficient in FXYD2 Regulatory Subunit of Na,K-ATPase. Front Physiol. 2016;7:88. doi:10.3389/fphys.2016.00088

52. Williams AL, Bohnsack BL. The Ocular Neural Crest: Specification, Migration, and Then What? Front Cell Dev Biol. 2020;8:595896. doi:10.3389/fcell.2020.595896

53. Cvekl A, Tamm ER. Anterior eye development and ocular mesenchyme: new insights from mouse models and human diseases. Bioessays. Apr 2004;26(4):374–386. doi:10.1002/bies.20009

54. Gong HY, Trinkaus-Randall V, Freddo TF. Ultrastructural immunocytochemical localization of elastin in normal human trabecular meshwork. Curr Eye Res. Oct 1989;8(10):1071–1082. doi:10.3109/02713688908997400

55. Xue W, Wallin R, Olmsted-Davis EA, Borrás T. Matrix GLA protein function in human trabecular meshwork cells: inhibition of BMP2-induced calcification process. Invest Ophthalmol Vis Sci. Mar 2006;47(3):997–1007. doi:10.1167/iovs.05-1106

56. Chowdhury UR, Jea SY, Oh DJ, Rhee DJ, Fautsch MP. Expression profile of the matricellular protein osteopontin in primary open-angle glaucoma and the normal human eye. Invest Ophthalmol Vis Sci. Aug 16 2011;52(9):6443–6451. doi:10.1167/iovs.11-7409

57. Young TL, Whisenhunt KN, Jin J, et al. SVEP1 as a Genetic Modifier of TEK-Related Primary Congenital Glaucoma. Invest Ophthalmol Vis Sci. Oct 1 2020;61(12):6. doi:10.1167/iovs.61.12.6

58. Narooie-Nejad M, Paylakhi SH, Shojaee S, et al. Loss of function mutations in the gene encoding latent transforming growth factor beta binding protein 2, LTBP2, cause primary congenital glaucoma. Hum Mol Genet. Oct 15 2009;18(20):3969–3977. doi:10.1093/hmg/ddp338

59. Lütjen-Drecoll E. Functional morphology of the trabecular meshwork in primate eyes. Prog Retin Eye Res. Jan 1999;18(1):91–119. doi:10.1016/s1350-9462(98)00011-1

60. Shuman MA, Polansky JR, Merkel C, Alvarado JA. Tissue plasminogen activator in cultured human trabecular meshwork cells. Predominance of enzyme over plasminogen activator inhibitor. Invest Ophthalmol Vis Sci. Mar 1988;29(3):401–405.

61. De La Paz MA, Epstein DL. Effect of age on superoxide dismutase activity of human trabecular meshwork. Invest Ophthalmol Vis Sci. Aug 1996;37(9):1849–1853.

62. Liton PB, Luna C, Bodman M, Hong A, Epstein DL, Gonzalez P. Induction of IL-6 expression by mechanical stress in the trabecular meshwork. Biochem Biophys Res Commun. Dec 2 2005;337(4):1229–1236. doi:10.1016/j.bbrc.2005.09.182

63. Rittig M, Flügel C, Prehm P, Lütjen-Drecoll E. Hyaluronan synthase immunoreactivity in the anterior segment of the primate eye. Graefes Arch Clin Exp Ophthalmol. Jun 1993;231(6):313–317. doi:10.1007/bf00919026

64. Usui T, Nakajima F, Ideta R, et al. Hyaluronan synthase in trabecular meshwork cells. Br J Ophthalmol. Mar 2003;87(3):357–360. doi:10.1136/bjo.87.3.357

65. Keller KE, Vranka JA, Haddadin RI, et al. The effects of tenascin C knockdown on trabecular meshwork outflow resistance. Invest Ophthalmol Vis Sci. Aug 19 2013;54(8):5613–5623. doi:10.1167/iovs.13-11620

66. Flügel-Koch C, Ohlmann A, Fuchshofer R, Welge-Lüssen U, Tamm ER. Thrombospondin-1 in the trabecular meshwork: localization in normal and glaucomatous eyes, and induction by TGF-beta1 and dexamethasone in vitro. Exp Eye Res. Nov 2004;79(5):649– 663. doi:10.1016/j.exer.2004.07.005

67. Siegner A, May CA, Welge-Lüssen UW, Bloemendal H, Lütjen-Drecoll E. alpha B-crystallin in the primate ciliary muscle and trabecular meshwork. Eur J Cell Biol. Oct 1996;71(2):165–169.

68. Walsh JG, Cullen SP, Sheridan C, Lüthi AU, Gerner C, Martin SJ. Executioner caspase-3 and caspase-7 are functionally distinct proteases. Proc Natl Acad Sci U S A. Sep 2 2008;105(35):12815–12819. doi:10.1073/pnas.0707715105

69. Matsumoto K, Shionyu M, Go M, et al. Distinct interaction of versican/PG-M with hyaluronan and link protein. J Biol Chem. Oct 17 2003;278(42):41205–41212. doi:10.1074/jbc.M305060200

70. Keller KE, Bradley JM, Vranka JA, Acott TS. Segmental versican expression in the trabecular meshwork and involvement in outflow facility. Invest Ophthalmol Vis Sci. Jul 7 2011;52(8):5049–5057. doi:10.1167/iovs.10-6948

71. McCulloch DR, Nelson CM, Dixon LJ, et al. ADAMTS metalloproteases generate active versican fragments that regulate interdigital web regression. Dev Cell. Nov 2009;17(5):687–698. doi:10.1016/j.devcel.2009.09.008

72. Dua HS, Faraj LA, Branch MJ, et al. The collagen matrix of the human trabecular meshwork is an extension of the novel pre-Descemet’s layer (Dua’s layer). Br J Ophthalmol. May 2014;98(5):691–697. doi:10.1136/bjophthalmol-2013-304593

73. Fang J, Hou F, Wu S, et al. Piezo2 downregulation via the Cre-lox system affects aqueous humor dynamics in mice. Mol Vis. 2021;27:354–364.

74. Docheva D, Hunziker EB, Fassler R, Brandau O. Tenomodulin is necessary for tenocyte proliferation and tendon maturation. Mol Cell Biol. Jan 2005;25(2):699–705. doi:10.1128/MCB.25.2.699-705.2005

75. Runager K, Enghild JJ, Klintworth GK. Focus on molecules: Transforming growth factor beta induced protein (TGFBIp). Exp Eye Res. Oct 2008;87(4):298–299. doi:10.1016/j.exer.2007.12.001

76. Harada K, Yamazaki T, Iwata C, et al. Identification of targets of Prox1 during in vitro vascular differentiation from embryonic stem cells: functional roles of HoxD8 in lymphangiogenesis. J Cell Sci. Nov 1 2009;122(Pt 21):3923–3930. doi:10.1242/jcs.052324

77. Ma AS, Grigg JR, Jamieson RV. Phenotype-genotype correlations and emerging pathways in ocular anterior segment dysgenesis. Hum Genet. Sep 2019;138(8-9):899–915. doi:10.1007/s00439-018-1935-7

78. Siggs OM, Souzeau E, Pasutto F, et al. Prevalence of FOXC1 Variants in Individuals With a Suspected Diagnosis of Primary Congenital Glaucoma. JAMA Ophthalmol. Apr 1 2019;137(4):348–355. doi:10.1001/jamaophthalmol.2018.5646

79. Siggs OM, Souzeau E, Taranath DA, et al. Biallelic CPAMD8 Variants Are a Frequent Cause of Childhood and Juvenile Open-Angle Glaucoma. Ophthalmology. Jun 2020;127(6):758–766. doi:10.1016/j.ophtha.2019.12.024

80. Tham YC, Li X, Wong TY, Quigley HA, Aung T, Cheng CY. Global prevalence of glaucoma and projections of glaucoma burden through 2040: a systematic review and meta-analysis. Ophthalmology. Nov 2014;121(11):2081–2090. doi:10.1016/j.ophtha.2014.05.013

81. Kwon YH, Fingert JH, Kuehn MH, Alward WL. Primary open-angle glaucoma. N Engl J Med. Mar 12 2009;360(11):1113–1124. doi:10.1056/NEJMra0804630

82. Gharahkhani P, Jorgenson E, Hysi P, et al. Genome-wide meta-analysis identifies 127 open-angle glaucoma loci with consistent effect across ancestries. Nat Commun. Feb 24 2021;12(1):1258. doi:10.1038/s41467-020-20851-4

83. Han X, Gharahkhani P, Hamel AR, et al. Large-scale multitrait genome-wide association analyses identify hundreds of glaucoma risk loci. Nat Genet. Jul 2023;55(7):1116–1125. doi:10.1038/s41588-023-01428-5

84. Zhao Y, Wang S, Sorenson CM, et al. Cyp1b1 mediates periostin regulation of trabecular meshwork development by suppression of oxidative stress. Mol Cell Biol. Nov 2013;33(21):4225–4240. doi:10.1128/mcb.00856-13

85. Yarishkin O, Phuong TTT, Baumann JM, et al. Piezo1 channels mediate trabecular meshwork mechanotransduction and promote aqueous fluid outflow. J Physiol. Jan 2021;599(2):571–592. doi:10.1113/JP281011

86. Du J, Thomson BR, Onay T, Quaggin SE. Endothelial Tyrosine Kinase Tie1 Is Required for Normal Schlemm’s Canal Development-Brief Report. Arterioscler Thromb Vasc Biol. Mar 2022;42(3):348–351. doi:10.1161/ATVBAHA.121.316692

87. Cheong SS, Hentschel L, Davidson AE, et al. Mutations in CPAMD8 Cause a Unique Form of Autosomal-Recessive Anterior Segment Dysgenesis. Am J Hum Genet. Dec 1 2016;99(6):1338–1352. doi:10.1016/j.ajhg.2016.09.022

88. Dulle JE, Rübsam A, Garnai SJ, Pawar HS, Fort PE. BetaB2-crystallin mutations associated with cataract and glaucoma leads to mitochondrial alterations in lens epithelial cells and retinal neurons. Exp Eye Res. Feb 2017;155:85–90. doi:10.1016/j.exer.2017.01.005

89. Garnai SJ, Huyghe JR, Reed DM, et al. Congenital cataracts: de novo gene conversion event in CRYBB2. Mol Vis. 2014;20:1579–1593.

90. Liu H, Bell K, Herrmann A, et al. Crystallins Play a Crucial Role in Glaucoma and Promote Neuronal Cell Survival in an In Vitro Model Through Modulating Müller Cell Secretion. Invest Ophthalmol Vis Sci. Jul 8 2022;63(8):3. doi:10.1167/iovs.63.8.3

91. Semina EV, Brownell I, Mintz-Hittner HA, Murray JC, Jamrich M. Mutations in the human forkhead transcription factor FOXE3 associated with anterior segment ocular dysgenesis and cataracts. Hum Mol Genet. Feb 1 2001;10(3):231–236. doi:10.1093/hmg/10.3.231

92. Blixt A, Mahlapuu M, Aitola M, Pelto-Huikko M, Enerbäck S, Carlsson P. A forkhead gene, FoxE3, is essential for lens epithelial proliferation and closure of the lens vesicle. Genes Dev. Jan 15 2000;14(2):245–254.

93. Semina EV, Ferrell RE, Mintz-Hittner HA, et al. A novel homeobox gene PITX3 is mutated in families with autosomal-dominant cataracts and ASMD. Nat Genet. Jun 1998;19(2):167–170. doi:10.1038/527

94. Nair KS, Hmani-Aifa M, Ali Z, et al. Alteration of the serine protease PRSS56 causes angle-closure glaucoma in mice and posterior microphthalmia in humans and mice. Nat Genet. Jun 2011;43(6):579–584. doi:10.1038/ng.813

95. Rad A, Altunoglu U, Miller R, et al. MAB21L1 loss of function causes a syndromic neurodevelopmental disorder with distinctive cerebellar, ocular, craniofacial and genital features (COFG syndrome). J Med Genet. May 2019;56(5):332–339. doi:10.1136/jmedgenet-2018-105623

96. Deml B, Kariminejad A, Borujerdi RH, Muheisen S, Reis LM, Semina EV. Mutations in MAB21L2 result in ocular Coloboma, microcornea and cataracts. PLoS Genet. 2015;11(2):e1005002. doi:10.1371/journal.pgen.1005002

97. Bruel AL, Masurel-Paulet A, Rivière JB, et al. Autosomal recessive truncating MAB21L1 mutation associated with a syndromic scrotal agenesis. Clin Genet. Feb 2017;91(2):333–338. doi:10.1111/cge.12794

98. Seese SE, Deml B, Muheisen S, Sorokina E, Semina EV. Genetic disruption of zebrafish mab21l1 reveals a conserved role in eye development and affected pathways. Dev Dyn. Aug 2021;250(8):1056–1073. doi:10.1002/dvdy.312

99. Wawersik S, Maas RL. Vertebrate eye development as modeled in Drosophila. Hum Mol Genet. Apr 12 2000;9(6):917–925. doi:10.1093/hmg/9.6.917

100. Hanson IM. PAX6 and congenital eye malformations. Pediatr Res. Dec 2003;54(6):791–796. doi:10.1203/01.Pdr.0000096455.00657.98

101. Baulmann DC, Ohlmann A, Flügel-Koch C, Goswami S, Cvekl A, Tamm ER. Pax6 heterozygous eyes show defects in chamber angle differentiation that are associated with a wide spectrum of other anterior eye segment abnormalities. Mech Dev. Oct 2002;118(1-2):3–17. doi:10.1016/s0925-4773(02)00260-5

102. Ferre-Fernández JJ, Aroca-Aguilar JD, Medina-Trillo C, et al. Whole-Exome Sequencing of Congenital Glaucoma Patients Reveals Hypermorphic Variants in GPATCH3, a New Gene Involved in Ocular and Craniofacial Development. Sci Rep. Apr 11 2017;7:46175. doi:10.1038/srep46175

103. Banerji S, Ni J, Wang SX, et al. LYVE-1, a new homologue of the CD44 glycoprotein, is a lymph-specific receptor for hyaluronan. J Cell Biol. Feb 22 1999;144(4):789–801. doi:10.1083/jcb.144.4.789

104. Smith T, Heger A, Sudbery I. UMI-tools: modeling sequencing errors in Unique Molecular Identifiers to improve quantification accuracy. Genome Res. Mar 2017;27(3):491–499. doi:10.1101/gr.209601.116

105. Hao Y, Stuart T, Kowalski MH, et al. Dictionary learning for integrative, multimodal and scalable single-cell analysis. Nat Biotechnol. Feb 2024;42(2):293–304. doi:10.1038/s41587-023-01767-y

106. Nguyen HCT, Baik B, Yoon S, Park T, Nam D. Benchmarking integration of single-cell differential expression. Nat Commun. Mar 21 2023;14(1):1570. doi:10.1038/s41467-023-37126-3

107. scCustomize: Custom Visualizations & Functions for Streamlined Analyses of Single Cell Sequencing. Version 2.1.2. Zenodo; 2024. https://samuel-marsh.github.io/scCustomize/

