## Supporting Information for "Leveraging single-cell transcriptomics of developing rat ocular outflow tissues to prioritize congenital glaucoma candidate genes"

### S1 Appendix

#### Identification of major limbal cell types

Data contributions from each of the 18 limbal samples (acquisition batches provided in Table S1) were largely consistent (4,020-6,171 cells (4.6-7.1% dataset)), except one that had a resuspension issue (P9 male, 1,595 cells (1.8% dataset); Table S2).

The identities of the 13 limbal cell clusters (C0-C12) were revealed through differential expression of published marker genes (Fig. 2). During the development of the eye, the periocular mesenchyme (POM) gives rise to several important structures, including the corneal stroma and endothelium, the iris stroma and muscles, the ciliary body stroma and muscles, portions of the sclera, the trabecular meshwork, and pericytes.[1,2] These tissues contributed to cell clusters C0, C1, C3 and C11, identified by their continued expression of the canonical POM marker *Pitx2*.[1,3] Smooth muscle-like cells formed C0 with high expression of *Pde5a*, *Tagln*, and *Acta2*.[4-6] Fibroblast-like cells, likely derived from the ocular connective tissues, such as the stroma of the cornea and sclera, and the TM, were located in C1 with higher expression of *Col1a1*, *Col6a3*, *Dcn*, *Adamts2*, and *Matn4*,[7-9] but lower expression of smooth muscle markers. Pericyte-like cells constituting C3 showed marked expression of *Rgs5*, *Kcnj8*, and *Vtn*.[10] Rapidly dividing mesenchymal progenitor-like cells in C11 were more prevalent during earlier stages of development and highly expressed fibroblast-like markers *Col1a1*, *Col6a3*, and *Dcn*, smooth muscle-like marker *Pde5a*, proliferation markers *Top2a*, *Prc1*, and *Racgap1*, and cell motility marker *Hmmr*.[11-14]

Known as the "master regulator" of eye development, *Pax6* is expressed in the iris, ciliary body and cornea,[15] and was restricted to clusters C2, C6 and C7. Pigmented epithelial cells from the iris and ciliary body formed C2, which expressed melanogenesis markers *Slc24a5*, *Mlana*, and *Tyr*.[16-18] Conversely, the closely related nonpigmented ciliary epithelium (NPCE) populated C6, which expressed markers *Col9a1*, *Lrp2*, and *Mecom*.[19-21] Finally, corneal, limbal, and conjunctival surface epithelium constituted C7, which highly expressed *Krt12* and *Krt15*.[22]

Melanocytes comprised C4, demonstrating the highest levels of the melanin synthesis markers *Slc24a5*, *Mlana*, and *Tyr*.[16-18] Monocytes expressing *Cd68* and *Cd14* formed C5 and C12, the latter differentiated via expression of anti-microbial receptor *Fpr2*.[23-25] Vascular endothelial cells strongly expressing canonical markers *Cdh5*, *Egfl7*, and *Pecam1* populated C8.[26-32] Schwann cells expressing *Cdh19* and *Lgi4* were represented in C9.[33,34] Finally, lymphocytes expressing *Icos* and *Cd69* occupied C10.[35,36]

#### Identification of VEC subtypes

The cell type identities within the VEC clusters were ascertained through interrogation of differentially expressed genes (Fig. S3). Blood vessel-like cells expressing *Cd34*, *Flt1*, and *Nos3* formed eight of the clusters (VEC0-7, Fig. 3).[27,29,37,38] Blood vessel markers for angiogenesis (*Sema3e*, *Stc1*, *Lrg1*),[39-41] as well as *Tcf15* and *Slc2a1* were enhanced in VEC0 and VEC3. Cells in VEC3 also exhibited markers of endothelial-to-mesenchymal transition (*Acta2*, *Pdgfrb*),[42,43] a process involved in the formation of vessel-associated connective tissues. Basement membrane expression (*Col4a5*) was elevated in VEC2.[44] Enhanced expression of an endothelial migration marker (*Id1*) was detected in VEC6 cells.[45] Cells expressing *Ackr1*, a marker of collector channels that link SC to the venous circulation, resided in VEC4 and VEC5.[46] Specifically, VEC4 cells expressed lower levels of *Ackr1* but higher markers of vessel tube formation (*Igf2*) and basement membrane deposition (*Lama3*), consistent with early collector channel development.[47,48] In contrast, lymphatic endothelial cells comprised the smaller VEC8 population with high expression of lymphatic markers *Prox1*, *Pdpn*, and *Lyve1,* and a lack of blood vascular markers (*Cd34*, *Flt1*, *Vwf*, *Nos3*).[29,49] SECs expressing hybrid features of blood and lymphatic endothelia (*Cd34* and *Prox1*, but not *Pdpn* or *Vwf*) formed VEC1 and a subpopulation of VEC7.[50-53] The two distinct VEC7 populations were formed from proliferating cells highly expressing *Top2a* and *Mki67*.[54,55] These populations also changed in size with tissue age (Fig. S3). As SC developed, the larger population displaying a stronger blood vessel-like profile (high *Flt1*, *Lrg1*, *Stc1*, *Sema3e*, *Tcf15*, low *Prox1*) reduced in number, whilst the smaller island exhibiting more SEC-like expression became numerous.

#### Identification of fibroblast-like subtypes

C1 cells were isolated and reclustered into four fibroblast-like (FBL0-FBL3) subtypes (Fig. S6A), and their identities inferred from established marker genes, as described in Supporting Information (Identification of FBL Subtypes, Fig. S6B and S7). In line with previous studies of adult human TM,[56] the pan-TM marker *Myoc* showed strongest expression in cluster FBL0, which also displayed high levels of other TM-associated genes including *Angpt1*,[57] *Eln*,[58] and *Thbs1*.[59] In contrast, FBL1 cells expressed markers of corneal stromal keratocytes, such as *Kera* and *Aldh3a1*,[60] while FBL2 cells were enriched for stromal markers *Cxcl14* and *Cd34*.[60,61] The small FBL3 satellite population expressed genes characteristic of corneal endothelium, including *Slc4a11*[62] and the basement membrane component *Col4a3*.[63]

#### S1 Appendix References


1. Williams AL, Bohnsack BL. The Ocular Neural Crest: Specification, Migration, and Then What? *Front Cell Dev Biol*. 2020;8:595896. doi:10.3389/fcell.2020.595896
2. Etchevers HC, Vincent C, Le Douarin NM, Couly GF. The cephalic neural crest provides pericytes and smooth muscle cells to all blood vessels of the face and forebrain. *Development*. Apr 2001;128(7):1059–1068. doi:10.1242/dev.128.7.1059
3. Hjalt TA, Semina EV, Amendt BA, Murray JC. The Pitx2 protein in mouse development. *Dev Dyn*. May 2000;218(1):195–200. doi:10.1002/(sici)1097-0177(200005)218:1<195::Aid-dvdy17>3.0.Co;2-c
4. Rybalkin SD, Rybalkina IG, Feil R, Hofmann F, Beavo JA. Regulation of cGMP-specific phosphodiesterase (PDE5) phosphorylation in smooth muscle cells. *J Biol Chem*. Feb 1 2002;277(5):3310–3317. doi:10.1074/jbc.M106562200
5. Tsuji-Tamura K, Morino-Koga S, Suzuki S, Ogawa M. The canonical smooth muscle cell marker TAGLN is present in endothelial cells and is involved in angiogenesis. *J Cell Sci*. Aug 1 2021;134(15)doi:10.1242/jcs.254920

6. McHugh KM, Crawford K, Lessard JL. A comprehensive analysis of the developmental and tissue-specific expression of the isoactin multigene family in the rat. *Dev Biol*. Dec 1991;148(2):442–458. doi:10.1016/0012-1606(91)90263-3
7. Muhl L, Genové G, Leptidis S, et al. Publisher Correction: Single-cell analysis uncovers fibroblast heterogeneity and criteria for fibroblast and mural cell identification and discrimination. *Nat Commun*. Sep 3 2020;11(1):4493. doi:10.1038/s41467-020-18511-8
8. Engel J, Furthmayr H, Odermatt E, et al. Structure and macromolecular organization of type VI collagen. *Ann N Y Acad Sci*. 1985;460:25–37. doi:10.1111/j.1749-6632.1985.tb51154.x
9. Klatt AR, Nitsche DP, Kobbe B, Macht M, Paulsson M, Wagener R. Molecular structure, processing, and tissue distribution of matrilin-4. *J Biol Chem*. May 18 2001;276(20):17267–17275. doi:10.1074/jbc.M100587200
10. van Splunder H, Villacampa P, Martínez-Romero A, Graupera M. Pericytes in the disease spotlight. *Trends Cell Biol*. Jan 2024;34(1):58–71. doi:10.1016/j.tcb.2023.06.001
11. Capranico G, Tinelli S, Austin CA, Fisher ML, Zunino F. Different patterns of gene expression of topoisomerase II isoforms in differentiated tissues during murine development. *Biochim Biophys Acta*. Aug 17 1992;1132(1):43–48. doi:10.1016/0167-4781(92)90050-a
12. Piunti A, Shilatifard A. The roles of Polycomb repressive complexes in mammalian development and cancer. *Nat Rev Mol Cell Biol*. May 2021;22(5):326–345. doi:10.1038/s41580-021-00341-1
13. Xu Z, Wu S, Tu J, et al. RACGAP1 promotes lung cancer cell proliferation through the PI3K/AKT signaling pathway. *Sci Rep*. Apr 15 2024;14(1):8694. doi:10.1038/s41598-024-58539-0
14. Savani RC, Wang C, Yang B, et al. Migration of bovine aortic smooth muscle cells after wounding injury. The role of hyaluronan and RHAMM. *J Clin Invest*. Mar 1995;95(3):1158–1168. doi:10.1172/jci117764
15. Shaham O, Menuchin Y, Farhy C, Ashery-Padan R. Pax6: a multi-level regulator of ocular development. *Prog Retin Eye Res*. Sep 2012;31(5):351–376. doi:10.1016/j.preteyeres.2012.04.002
16. Lamason RL, Mohideen MA, Mest JR, et al. SLC24A5, a putative cation exchanger, affects pigmentation in zebrafish and humans. *Science*. Dec 16 2005;310(5755):1782–1786. doi:10.1126/science.1116238
17. De Mazière AM, Muehlethaler K, van Donselaar E, et al. The melanocytic protein Melan-A/MART-1 has a subcellular localization distinct from typical melanosomal proteins. *Traffic*. Sep 2002;3(9):678–693. doi:10.1034/j.1600-0854.2002.30909.x
18. Lerner AB, Fitzpatrick TB, et al. Mammalian tyrosinase; preparation and properties. *J Biol Chem*. Mar 1949;178(1):185–195.
19. Liu CY, Olsen BR, Kao WW. Developmental patterns of two alpha 1(IX) collagen mRNA isoforms in mouse. *Dev Dyn*. Oct 1993;198(2):150–157. doi:10.1002/aja.1001980208
20. Storm T, Heegaard S, Christensen EI, Nielsen R. Megalin-deficiency causes high myopia, retinal pigment epithelium-macromelanosomes and abnormal development of the ciliary body in mice. *Cell Tissue Res*. Oct 2014;358(1):99–107. doi:10.1007/s00441-014-1919-4
21. van Zyl T, Yan W, McAdams AM, Monavarfeshani A, Hageman GS, Sanes JR. Cell atlas of the human ocular anterior segment: Tissue-specific and shared cell types. *Proc Natl Acad Sci U S A*. Jul 19 2022;119(29):e2200914119. doi:10.1073/pnas.2200914119
22. Yoshida S, Shimmura S, Kawakita T, et al. Cytokeratin 15 can be used to identify the limbal phenotype in normal and diseased ocular surfaces. *Invest Ophthalmol Vis Sci*. Nov 2006;47(11):4780–4786. doi:10.1167/iovs.06-0574
23. Choudhary M, Malek G. CD68: Potential Contributor to Inflammation and RPE Cell Dystrophy. *Adv Exp Med Biol*. 2023;1415:207–213. doi:10.1007/978-3-031-27681-1_30
24. Sharygin D, Koniaris LG, Wells C, Zimmers TA, Hamidi T. Role of CD14 in human disease. *Immunology*. Jul 2023;169(3):260–270. doi:10.1111/imm.13634
25. Le Y, Murphy PM, Wang JM. Formyl-peptide receptors revisited. *Trends Immunol*. Nov 2002;23(11):541–548. doi:10.1016/s1471-4906(02)02316-5
26. Newman PJ. The role of PECAM-1 in vascular cell biology. *Ann N Y Acad Sci*. Apr 18 1994;714:165–174. doi:10.1111/j.1749-6632.1994.tb12041.x

27. Pusztaszeri MP, Seelentag W, Bosman FT. Immunohistochemical expression of endothelial markers CD31, CD34, von Willebrand factor, and Fli-1 in normal human tissues. *J Histochem Cytochem*. Apr 2006;54(4):385–395. doi:10.1369/jhc.4A6514.2005
28. Schmidt D, von Hochstetter AR. The use of CD31 and collagen IV as vascular markers. A study of 56 vascular lesions. *Pathol Res Pract*. Jun 1995;191(5):410–414. doi:10.1016/s0344-0338(11)80727-2
29. Baluk P, McDonald DM. Markers for microscopic imaging of lymphangiogenesis and angiogenesis. *Ann N Y Acad Sci*. 2008;1131:1–12. doi:10.1196/annals.1413.001
30. Lampugnani MG, Resnati M, Raiteri M, et al. A novel endothelial-specific membrane protein is a marker of cell-cell contacts. *J Cell Biol*. Sep 1992;118(6):1511–1522. doi:10.1083/jcb.118.6.1511
31. Dejana E, Bazzoni G, Lampugnani MG. Vascular endothelial (VE)-cadherin: only an intercellular glue? *Exp Cell Res*. Oct 10 1999;252(1):13–19. doi:10.1006/excr.1999.4601
32. Nichol D, Stuhlmann H. EGFL7: a unique angiogenic signaling factor in vascular development and disease. *Blood*. Feb 9 2012;119(6):1345–1352. doi:10.1182/blood-2011-10-322446
33. Takahashi M, Osumi N. Identification of a novel type II classical cadherin: rat cadherin19 is expressed in the cranial ganglia and Schwann cell precursors during development. *Dev Dyn*. Jan 2005;232(1):200–208. doi:10.1002/dvdy.20209
34. Bermingham JR, Jr., Shearin H, Pennington J, et al. The claw paw mutation reveals a role for Lgi4 in peripheral nerve development. *Nat Neurosci*. Jan 2006;9(1):76–84. doi:10.1038/nn1598
35. Hutloff A, Dittrich AM, Beier KC, et al. ICOS is an inducible T-cell co-stimulator structurally and functionally related to CD28. *Nature*. Jan 21 1999;397(6716):263–266. doi:10.1038/16717
36. López-Cabrera M, Santis AG, Fernández-Ruiz E, et al. Molecular cloning, expression, and chromosomal localization of the human earliest lymphocyte activation antigen AIM/CD69, a new member of the C-type animal lectin superfamily of signal-transmitting receptors. *J Exp Med*. Aug 1 1993;178(2):537–547. doi:10.1084/jem.178.2.537
37. Shibuya M. Differential roles of vascular endothelial growth factor receptor-1 and receptor-2 in angiogenesis. *J Biochem Mol Biol*. Sep 30 2006;39(5):469–478. doi:10.5483/bmbrep.2006.39.5.469
38. Pollock JS, Nakane M, Buttery LD, et al. Characterization and localization of endothelial nitric oxide synthase using specific monoclonal antibodies. *Am J Physiol*. Nov 1993;265(5 Pt 1):C1379–1387. doi:10.1152/ajpcell.1993.265.5.C1379
39. Gu C, Yoshida Y, Livet J, et al. Semaphorin 3E and plexin-D1 control vascular pattern independently of neuropilins. *Science*. Jan 14 2005;307(5707):265–268. doi:10.1126/science.1105416
40. Sheikh-Hamad D. Mammalian stanniocalcin-1 activates mitochondrial antioxidant pathways: new paradigms for regulation of macrophages and endothelium. *Am J Physiol Renal Physiol*. Feb 2010;298(2):F248–254. doi:10.1152/ajprenal.00260.2009
41. Wang X, Abraham S, McKenzie JAG, et al. LRG1 promotes angiogenesis by modulating endothelial TGF-β signalling. *Nature*. Jul 18 2013;499(7458):306–311. doi:10.1038/nature12345
42. Alharthi A, Verma A, Sabbineni H, Adil MS, Somanath PR. Distinct effects of pharmacological inhibition of stromelysin1 on endothelial-to-mesenchymal transition and myofibroblast differentiation. *J Cell Physiol*. Jul 2021;236(7):5147–5161. doi:10.1002/jcp.30221
43. Hellström M, Kalén M, Lindahl P, Abramsson A, Betsholtz C. Role of PDGF-B and PDGFR-beta in recruitment of vascular smooth muscle cells and pericytes during embryonic blood vessel formation in the mouse. *Development*. Jun 1999;126(14):3047–3055. doi:10.1242/dev.126.14.3047
44. Sund M, Maeshima Y, Kalluri R. Bifunctional promoter of type IV collagen COL4A5 and COL4A6 genes regulates the expression of alpha5 and alpha6 chains in a distinct cell-specific fashion. *Biochem J*. May 1 2005;387(Pt 3):755–761. doi:10.1042/bj20041870
45. Ling F, Kang B, Sun XH. Id proteins: small molecules, mighty regulators. *Curr Top Dev Biol*. 2014;110:189–216. doi:10.1016/b978-0-12-405943-6.00005-1
46. van Zyl T, Yan W, McAdams A, et al. Cell atlas of aqueous humor outflow pathways in eyes of humans and four model species provides insight into glaucoma pathogenesis. *Proc Natl Acad Sci U S A*. May 12 2020;117(19):10339–10349. doi:10.1073/pnas.2001250117

47. Bach LA. Endothelial cells and the IGF system. *J Mol Endocrinol*. Feb 2015;54(1):R1–13. doi:10.1530/jme-14-0215
48. Abrass CK, Berfield AK, Ryan MC, Carter WG, Hansen KM. Abnormal development of glomerular endothelial and mesangial cells in mice with targeted disruption of the lama3 gene. *Kidney Int*. Sep 2006;70(6):1062–1071. doi:10.1038/sj.ki.5001706
49. Cossutta M, Darche M, Carpentier G, et al. Weibel-Palade Bodies Orchestrate Pericytes During Angiogenesis. *Arterioscler Thromb Vasc Biol*. Sep 2019;39(9):1843–1858. doi:10.1161/atvbaha.119.313021
50. Kizhatil K, Ryan M, Marchant JK, Henrich S, John SW. Schlemm's canal is a unique vessel with a combination of blood vascular and lymphatic phenotypes that forms by a novel developmental process. *PLoS Biol*. Jul 2014;12(7):e1001912. doi:10.1371/journal.pbio.1001912
51. Park DY, Lee J, Park I, et al. Lymphatic regulator PROX1 determines Schlemm's canal integrity and identity. *J Clin Invest*. Sep 2014;124(9):3960–3974. doi:10.1172/jci75392
52. Aspelund A, Tammela T, Antila S, et al. The Schlemm's canal is a VEGF-C/VEGFR-3-responsive lymphatic-like vessel. *J Clin Invest*. Sep 2014;124(9):3975–3986. doi:10.1172/jci75395
53. Balasubramanian R, Kizhatil K, Li T, et al. Transcriptomic profiling of Schlemm's canal cells reveals a lymphatic-biased identity and three major cell states. *Elife*. Oct 18 2024;13doi:10.7554/eLife.96459
54. Lang AJ, Mirski SE, Cummings HJ, Yu Q, Gerlach JH, Cole SP. Structural organization of the human TOP2A and TOP2B genes. *Gene*. Oct 23 1998;221(2):255–266. doi:10.1016/s0378-1119(98)00468-5
55. Gerdes J, Lemke H, Baisch H, Wacker HH, Schwab U, Stein H. Cell cycle analysis of a cell proliferation-associated human nuclear antigen defined by the monoclonal antibody Ki-67. *J Immunol*. Oct 1984;133(4):1710–1715.
56. Patel G, Fury W, Yang H, et al. Molecular taxonomy of human ocular outflow tissues defined by single-cell transcriptomics. *Proc Natl Acad Sci U S A*. Jun 9 2020;117(23):12856–12867. doi:10.1073/pnas.2001896117
57. Thomson BR, Liu P, Onay T, et al. Cellular crosstalk regulates the aqueous humor outflow pathway and provides new targets for glaucoma therapies. *Nat Commun*. Oct 18 2021;12(1):6072. doi:10.1038/s41467-021-26346-0
58. Hann CR, Fautsch MP. The elastin fiber system between and adjacent to collector channels in the human juxtacanalicular tissue. *Invest Ophthalmol Vis Sci*. Jan 2011;52(1):45–50. doi:10.1167/iovs.10-5620
59. Flügel-Koch C, Ohlmann A, Fuchshofer R, Welge-Lüssen U, Tamm ER. Thrombospondin-1 in the trabecular meshwork: localization in normal and glaucomatous eyes, and induction by TGF-beta1 and dexamethasone in vitro. *Exp Eye Res*. Nov 2004;79(5):649–663. doi:10.1016/j.exer.2004.07.005
60. Català P, Groen N, Dehnen JA, et al. Single cell transcriptomics reveals the heterogeneity of the human cornea to identify novel markers of the limbus and stroma. *Sci Rep*. Nov 5 2021;11(1):21727. doi:10.1038/s41598-021-01015-w
61. Liu L, Yu Y, Peng Q, et al. Distribution of Stromal Cell Subsets in Cultures from Distinct Ocular Surface Compartments. *J Ophthalmic Vis Res*. Oct–Dec 2020;15(4):493–501. doi:10.18502/jovr.v15i4.7780
62. Malhotra D, Loganathan SK, Chiu AM, Lukowski CM, Casey JR. Human Corneal Expression of SLC4A11, a Gene Mutated in Endothelial Corneal Dystrophies. *Sci Rep*. Jul 4 2019;9(1):9681. doi:10.1038/s41598-019-46094-y
63. Yellore VS, Rayner SA, Nguyen CK, et al. Analysis of the role of ZEB1 in the pathogenesis of posterior polymorphous corneal dystrophy. *Invest Ophthalmol Vis Sci*. Jan 25 2012;53(1):273–278. doi:10.1167/iovs.11-8038

### Supporting Figures

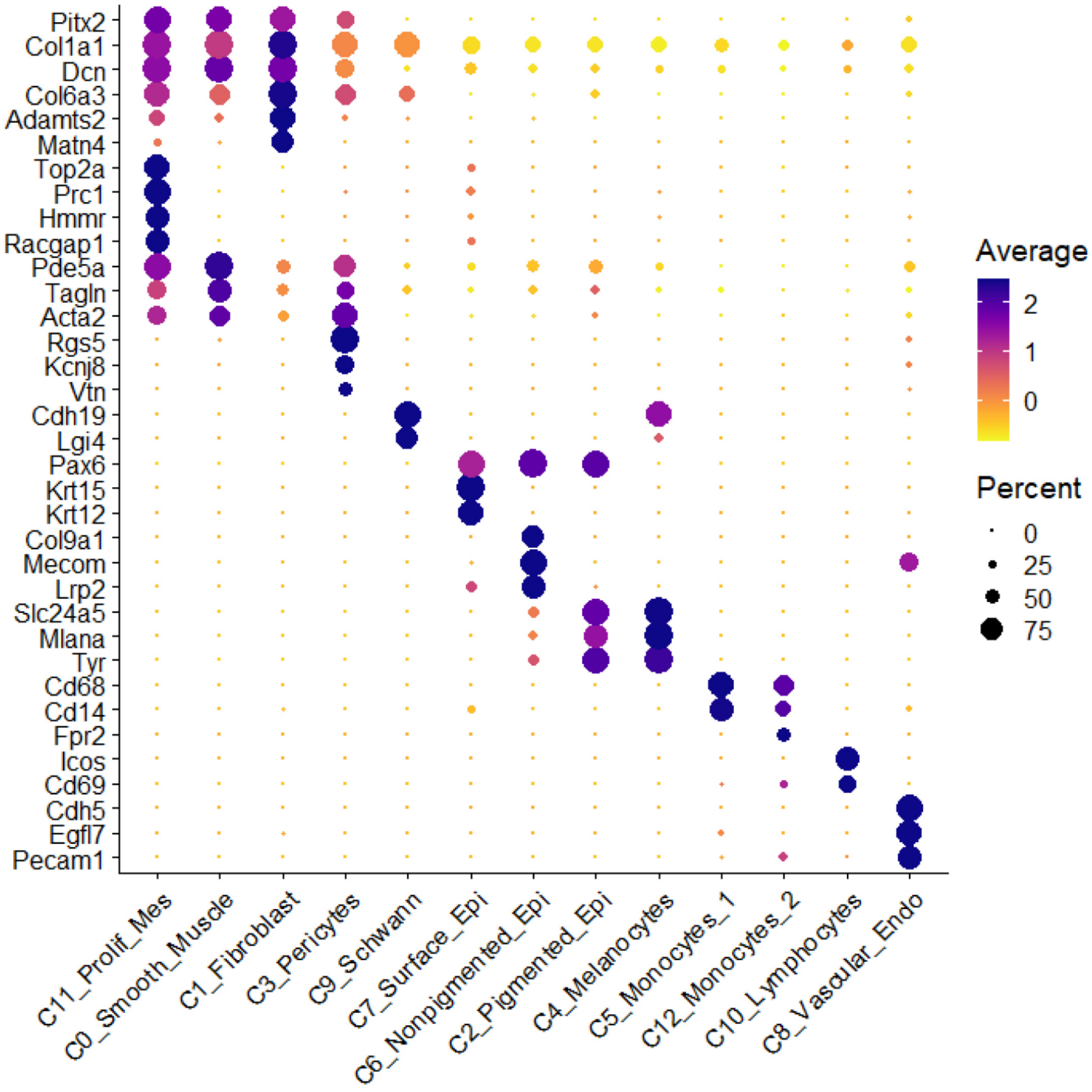


**Figure S1: Major limbal cell type clusters identified by key marker gene expression.** Dot plot showing selective expression of key marker genes that revealed cell type identities. The dot diameter represents the percentage of each cell type expressing the gene, while the color denotes the scaled average expression within the cluster. Epi, epithelium; Endo, endothelium; Prolif_Mes, proliferating mesenchyme.

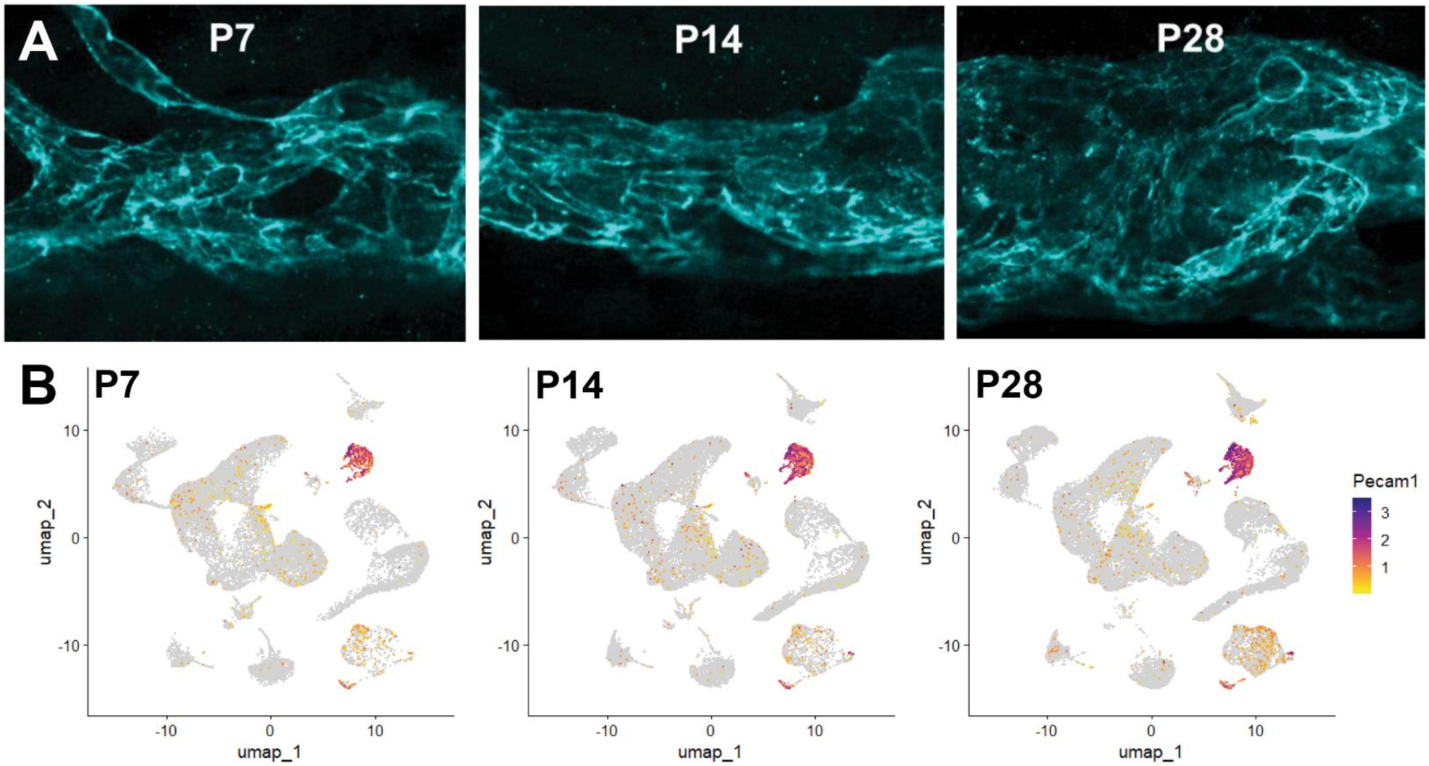


**Figure S2: Expression of Pecam1 identified vascular endothelial cell cluster containing SC cells.** (A) Immunofluorescent confocal imaging of developing SC in rat whole-mount anterior segments stained for Pecam1. At P7, SC is still developing with regions of smaller unfused vessels. At P14, most vessels have remodeled into a single continuous canal. At P28, SC has matured and grown significantly in diameter. (B) UMAP plot of clustered limbal cells with expression of Pecam1 mRNA at P7 (left), P14 (center), and P28 (right). Pecam1 expression is shown normalized on a log2 scale.

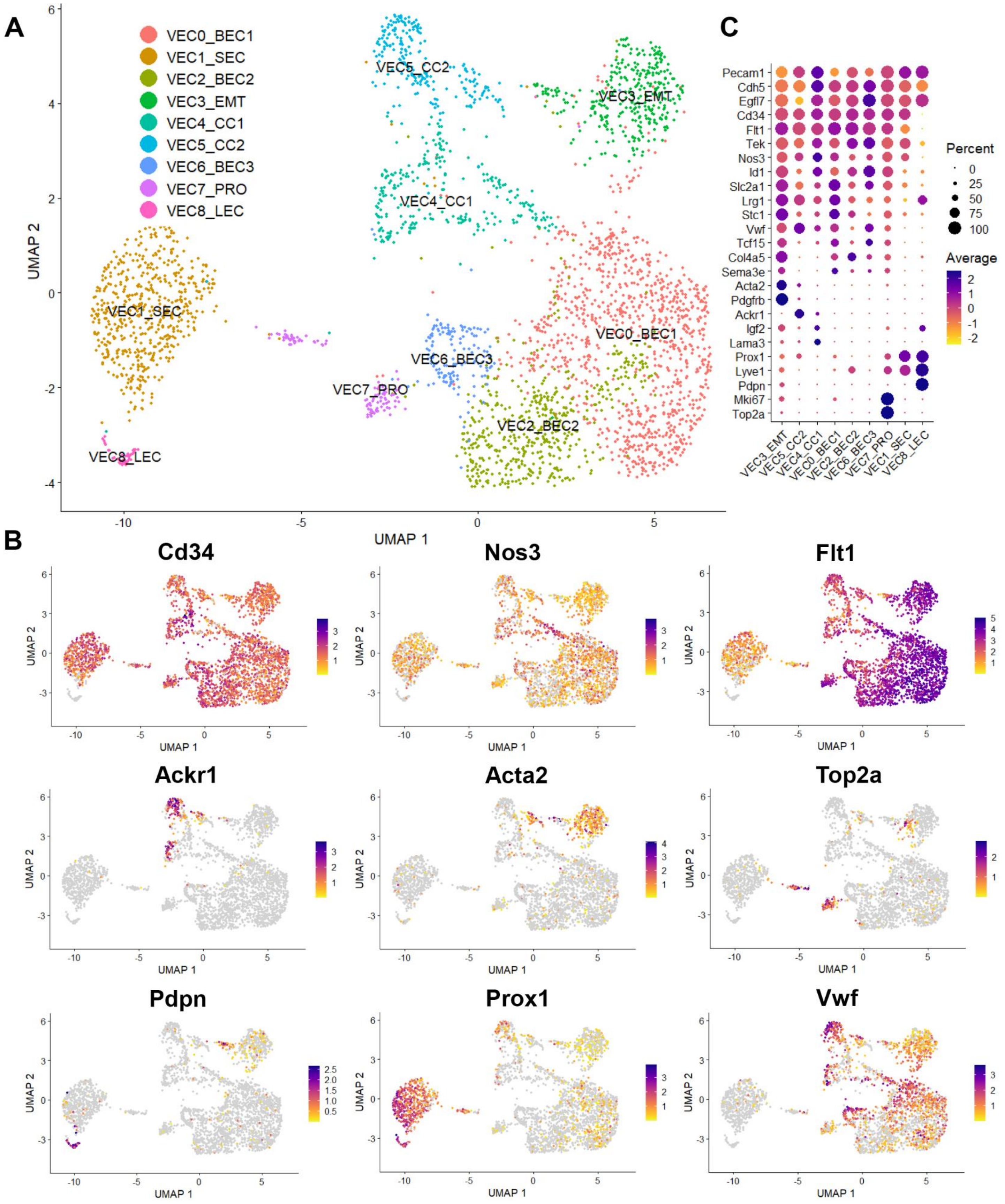


**Figure S3: Identification of Schlemm's canal endothelial cells amongst other vascular endothelial cell subtypes.** (A) Reclustering of VECs from the limbal C8 population visualized by UMAP in nine subtypes (VEC0-VEC8). (B) Normalized log2 fold change expression of 9 key VEC markers revealed cluster identities.

(C) Dot plot summarizing the scaled average expression of key VEC markers and the percentage of expressing cells within each of the 9 VEC subcluster types. BEC, blood VECs; SEC, Schlemm's canal VECs; EMT, VECs undergoing endothelial-to-mesenchymal transition; CC, collector channel VECs; PRO, proliferating VECs; LEC, lymphatic VECs.

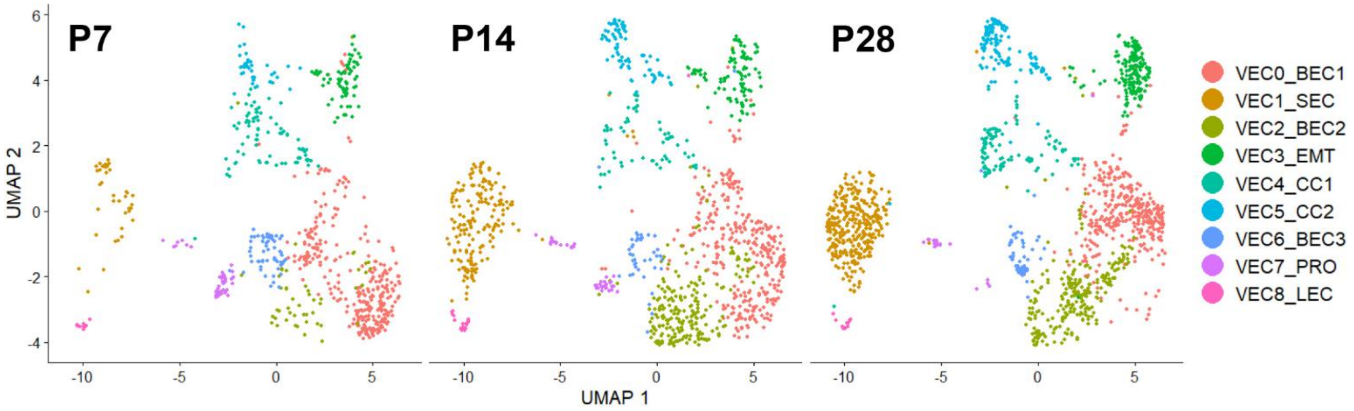


**Figure S4: Increase in VEC1 population across developmental age correlates with growth and maturation of Schlemm's canal.** UMAP plot of VEC subtype populations at each of the three developmental timepoints. Across the period that SC is developing, cells within the putative SC cluster (VEC1) show the most significant population increase, whilst the cell numbers within the proliferating VEC population (VEC7) diminish.

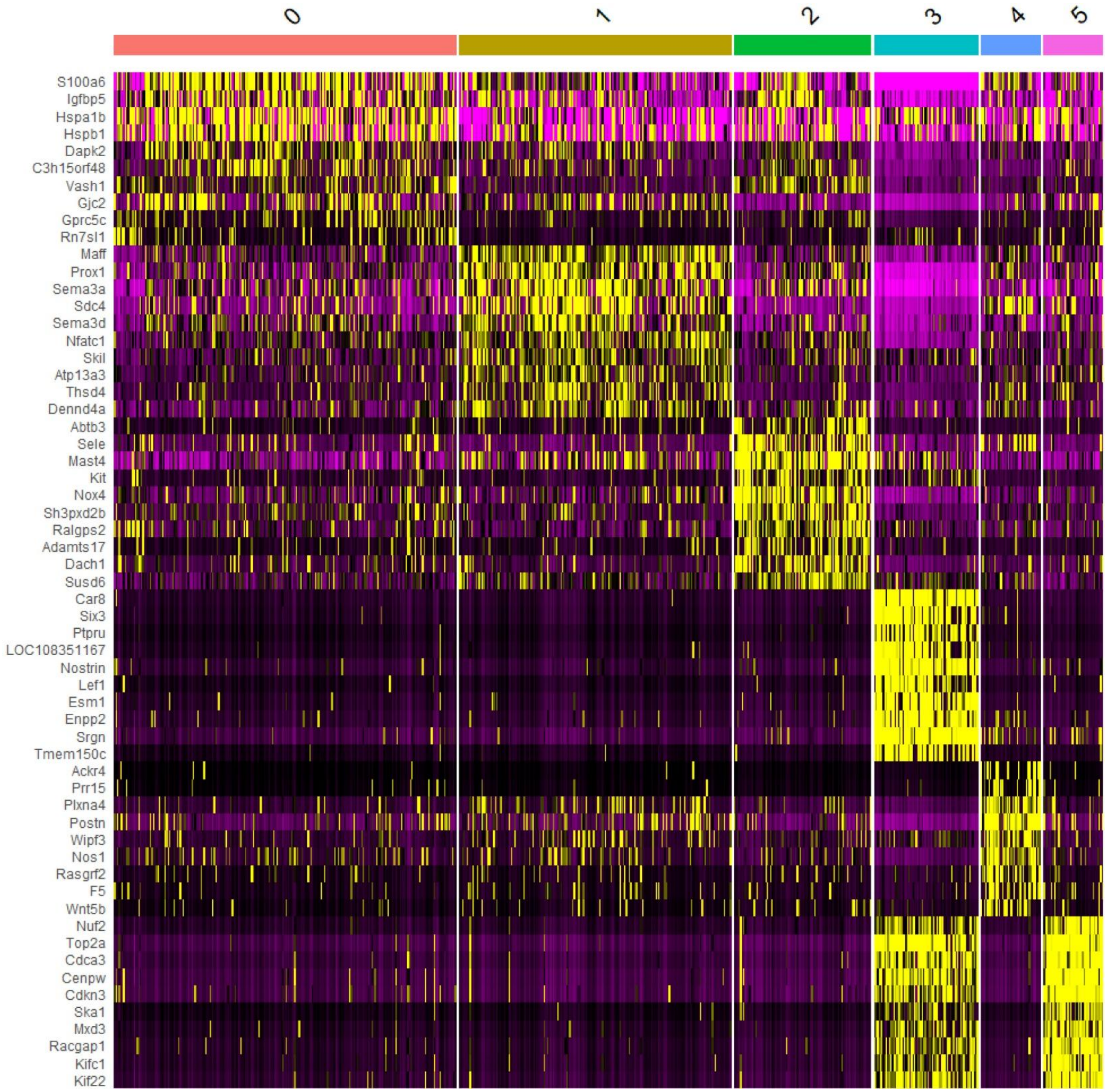


**Figure S5: Heat map illustrating the top 10 differentially expressed genes associated with each of the six SEC subtypes.** Clusters are shown as columns with specific genes as rows. For every cell (vertical line), the average log 2 fold-change in gene expression is depicted as upregulated (yellow) or downregulated (purple).

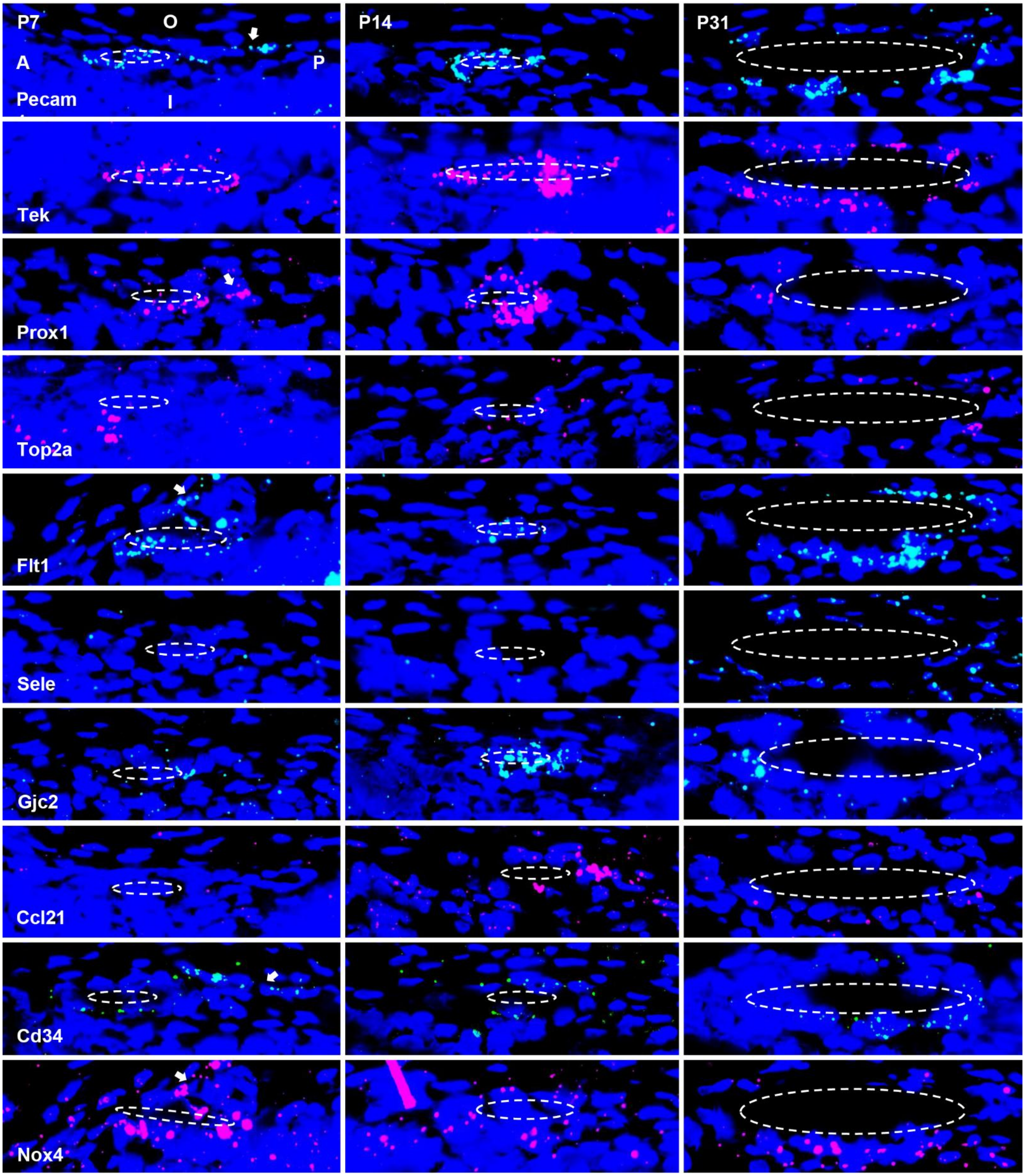


**Figure S6: Tissue localization of SC-associated marker gene expression.** Tissue sections (5µm) stained by *in situ* hybridization (ISH) for *Pecam1*, *Tek*, *Prox1*, *Top2a*, *Flt1*, *Sele*, *Gjc2*, *Ccl21*, *Cd34*, and *Nox4* transcripts. Nuclei counterstained with DAPI (blue). Representative tissue sections at P7 (left), P14 (center), and P31 (right). All images at same scale and oriented as shown in top-left image. A, anterior; P, posterior; O, outer side; I, inner side. SC locations (dashed ellipses) drawn from corresponding autofluorescence images. Presence of collector channels noted (arrows).

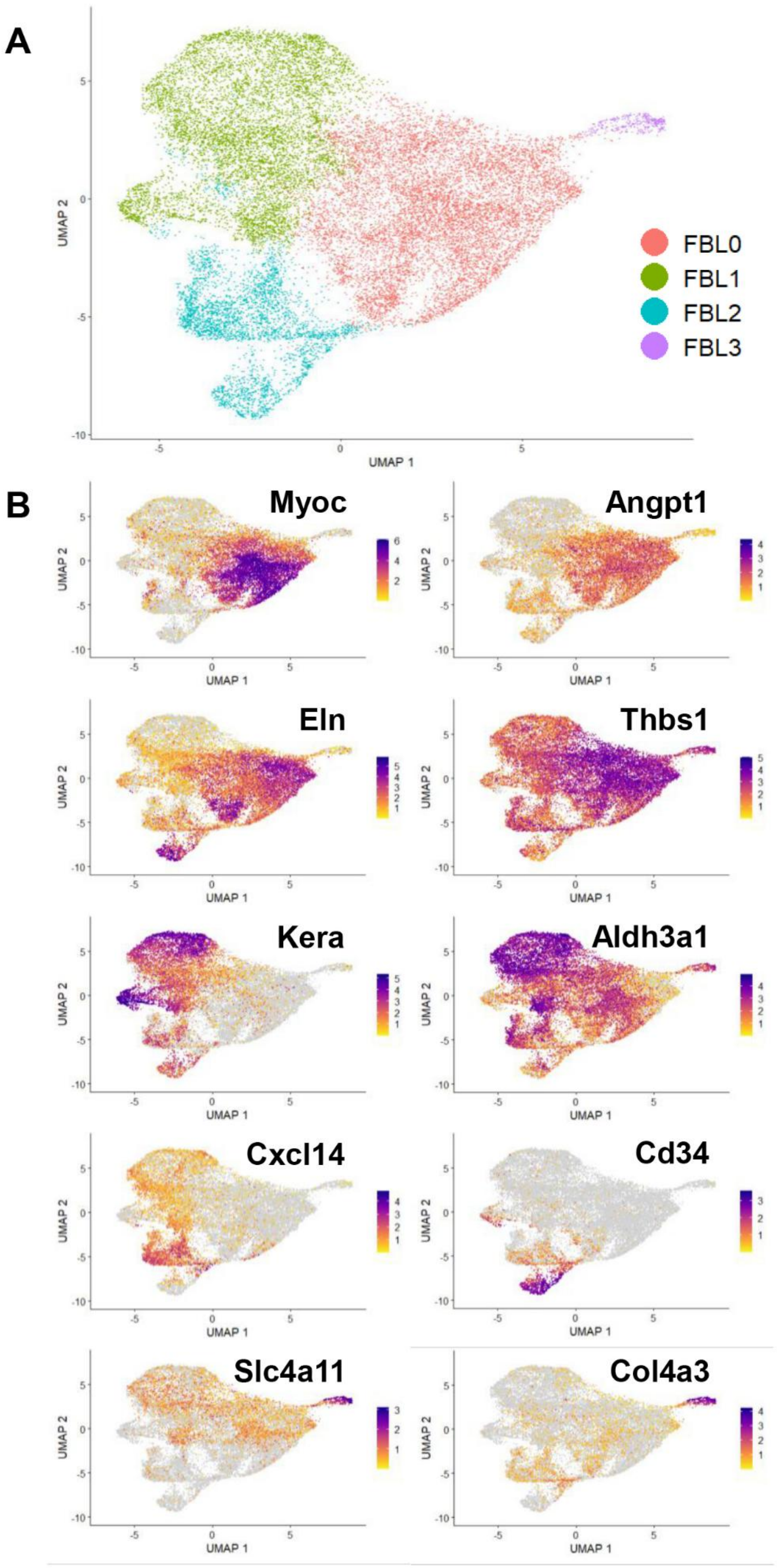


**Figure S7: FBL cluster identities inferred from established marker gene expression.** (A) UMAP representing C1 cells reclustered into four fibroblast-like (FBL) subtypes. (B) Expression of key markers (normalized log2 fold change) distinguishing cluster identities.

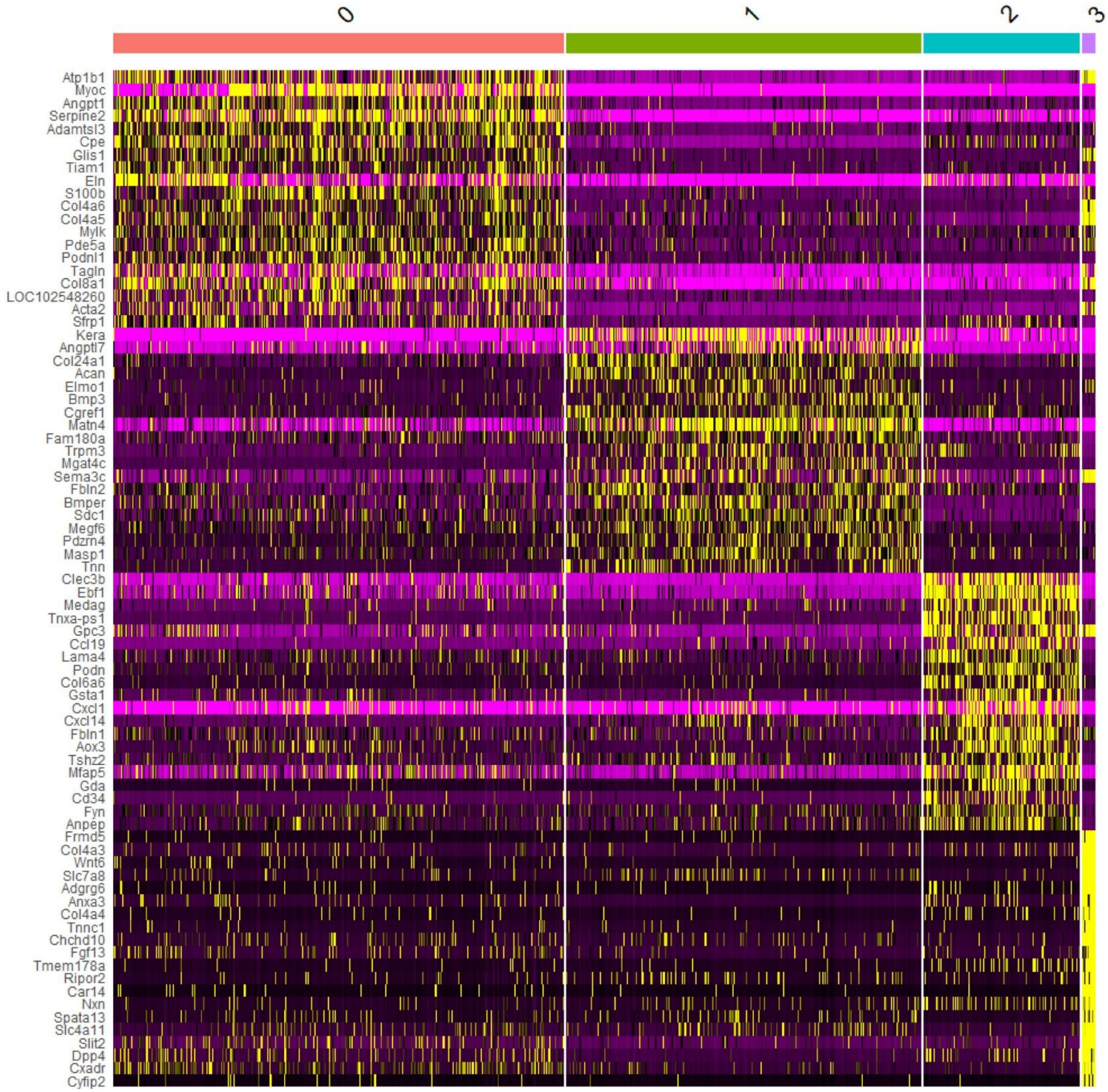


**Figure S8: Heat map showing the top 20 genes differentially expressed between the four fibroblast-like (FBL0-FBL3) subtypes within limbal cluster C1.**

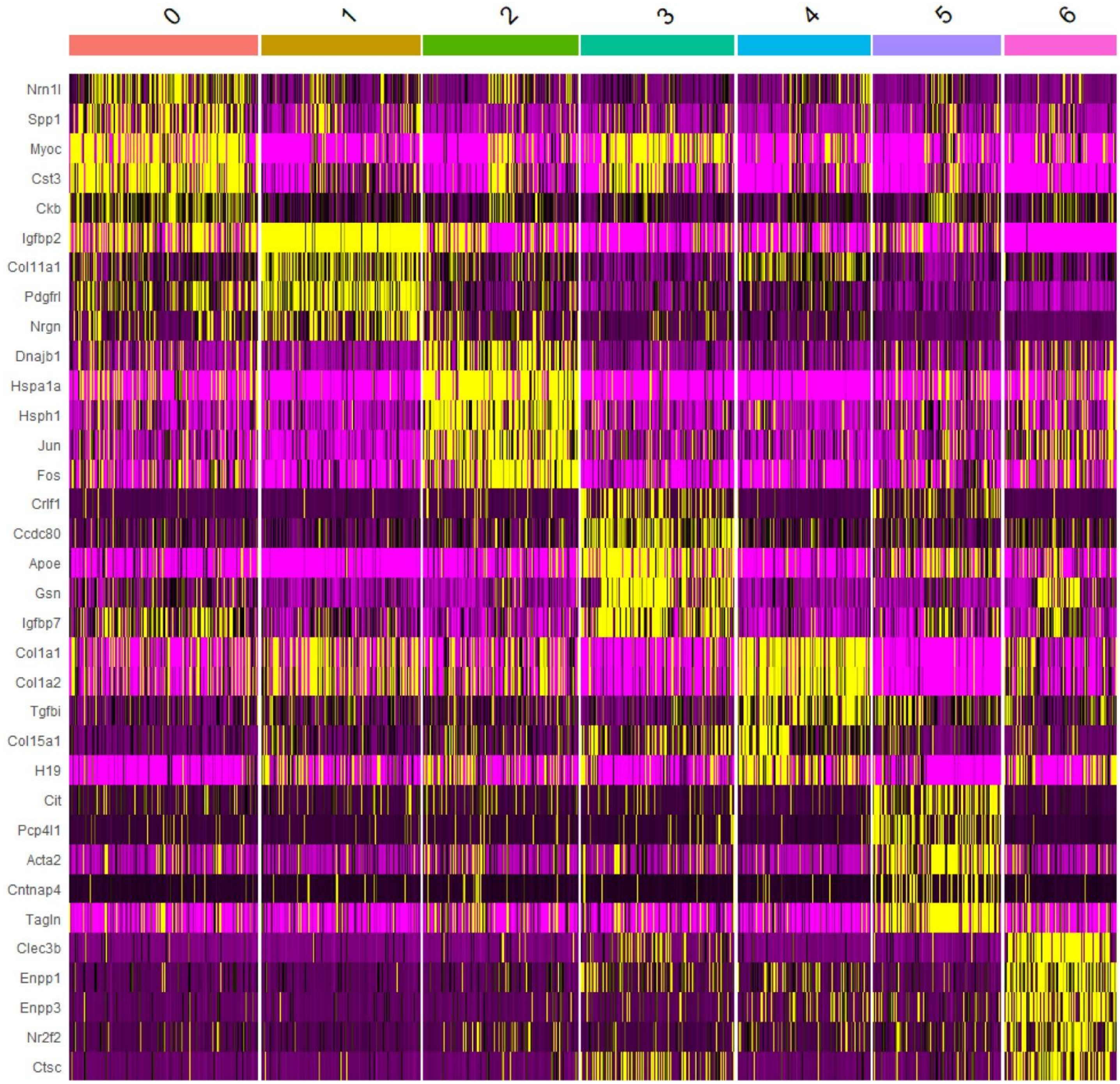


**Figure S9: Fig. Heat map showing up to top 5 genes differentially expressed between the seven TMC-like clusters (TMC0-TMC6) derived from fibroblast-like cluster FBL0.**

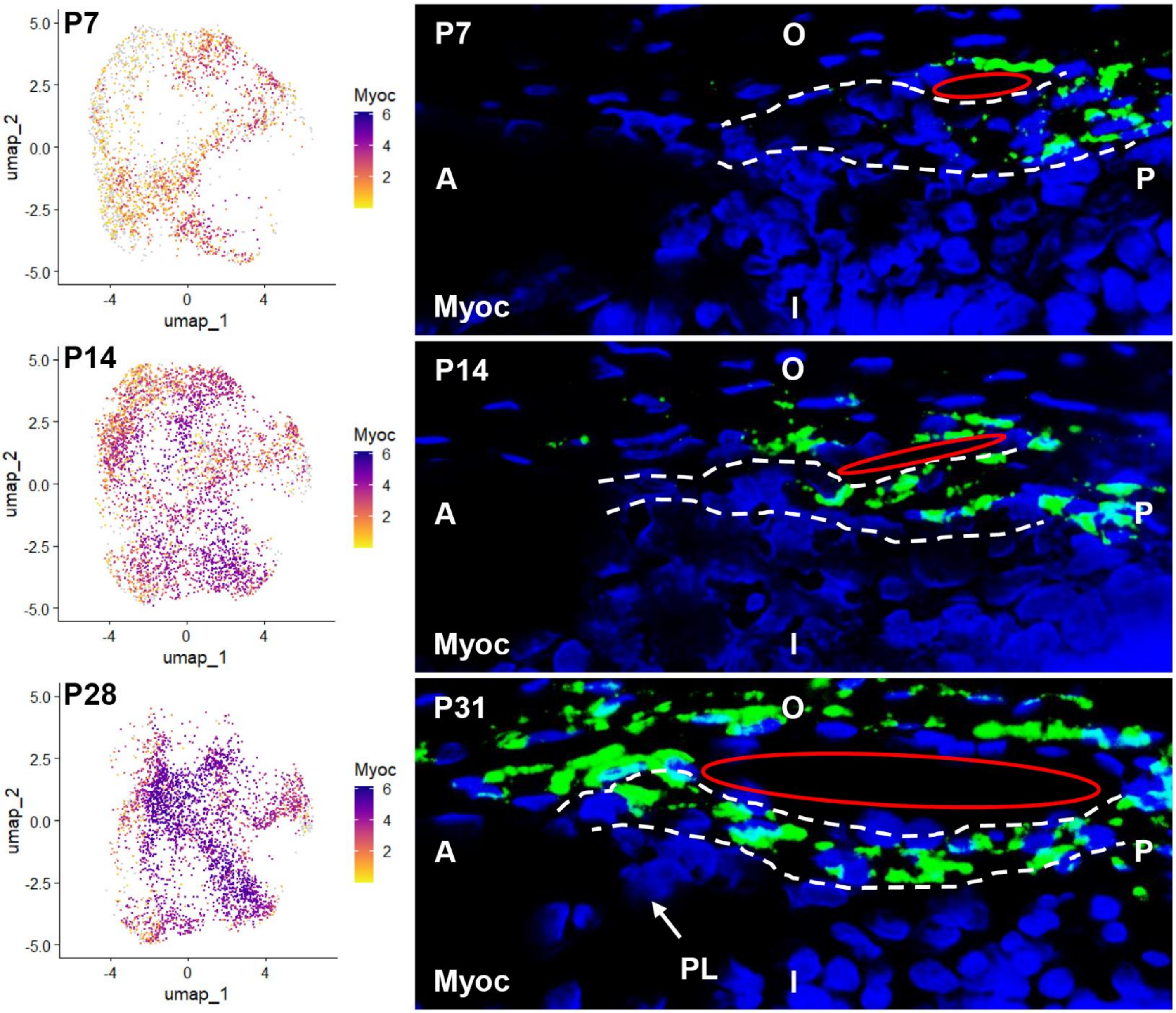


**Figure S10: Expression of *Myoc* within TMCs.** Expression for *Myoc* is shown in the scRNAseq data (normalized log2 fold change, left) alongside 5µm tissue sections stained by ISH (right) for each developmental age.

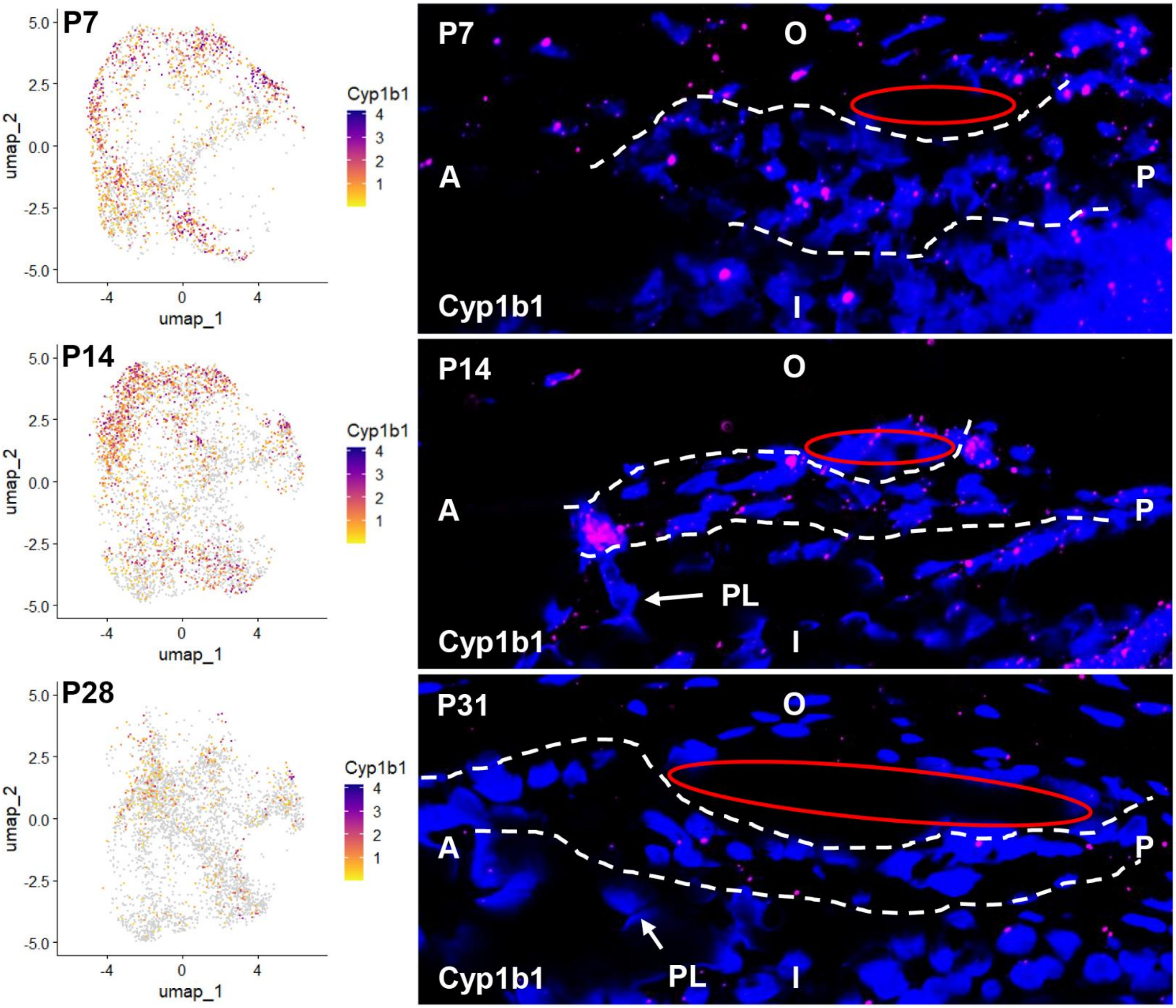


**Figure S11: Expression of *Cyp1b1* within TMCs.**

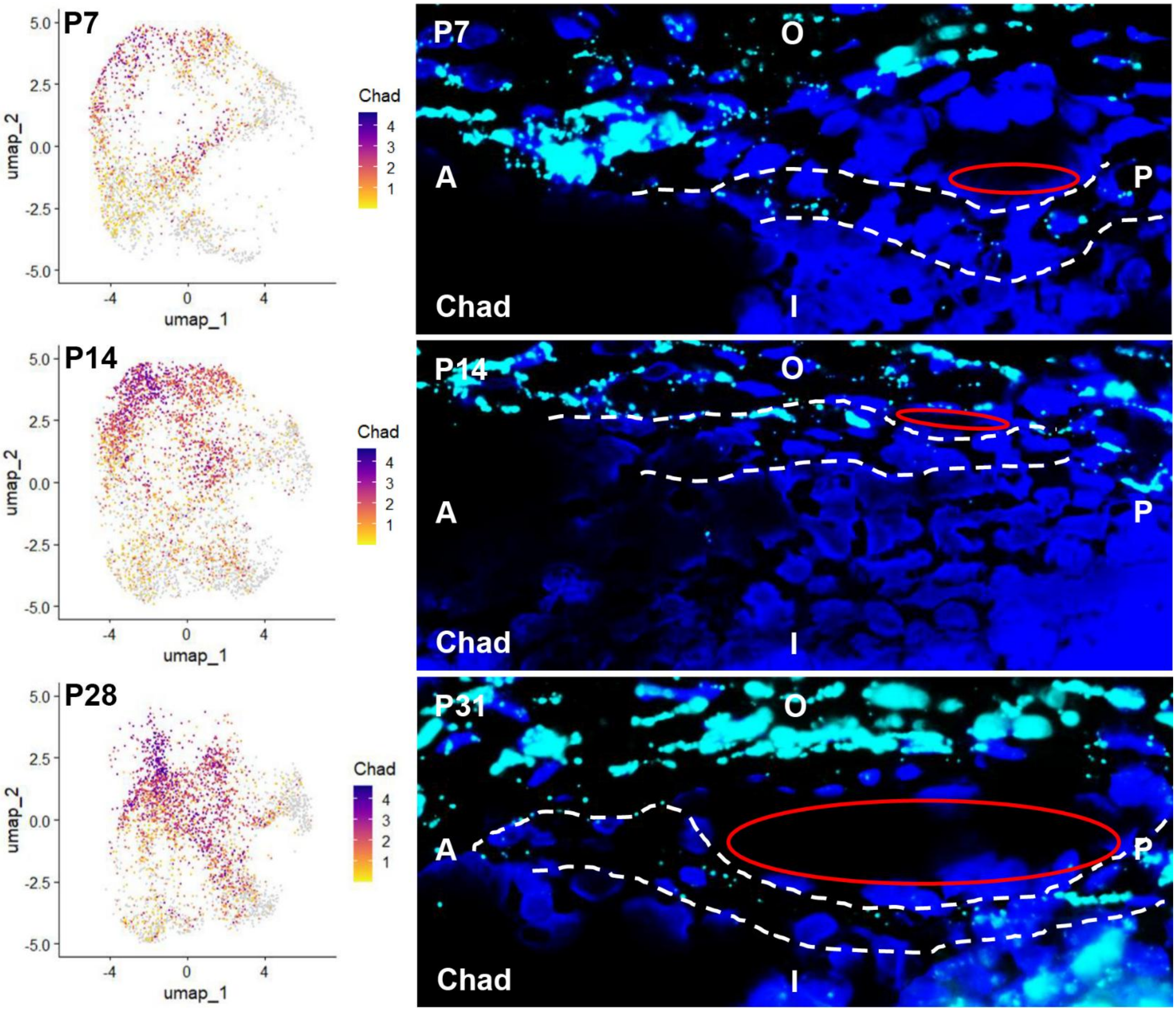

**Figure S12: Expression of *Chad* within TMCs.**

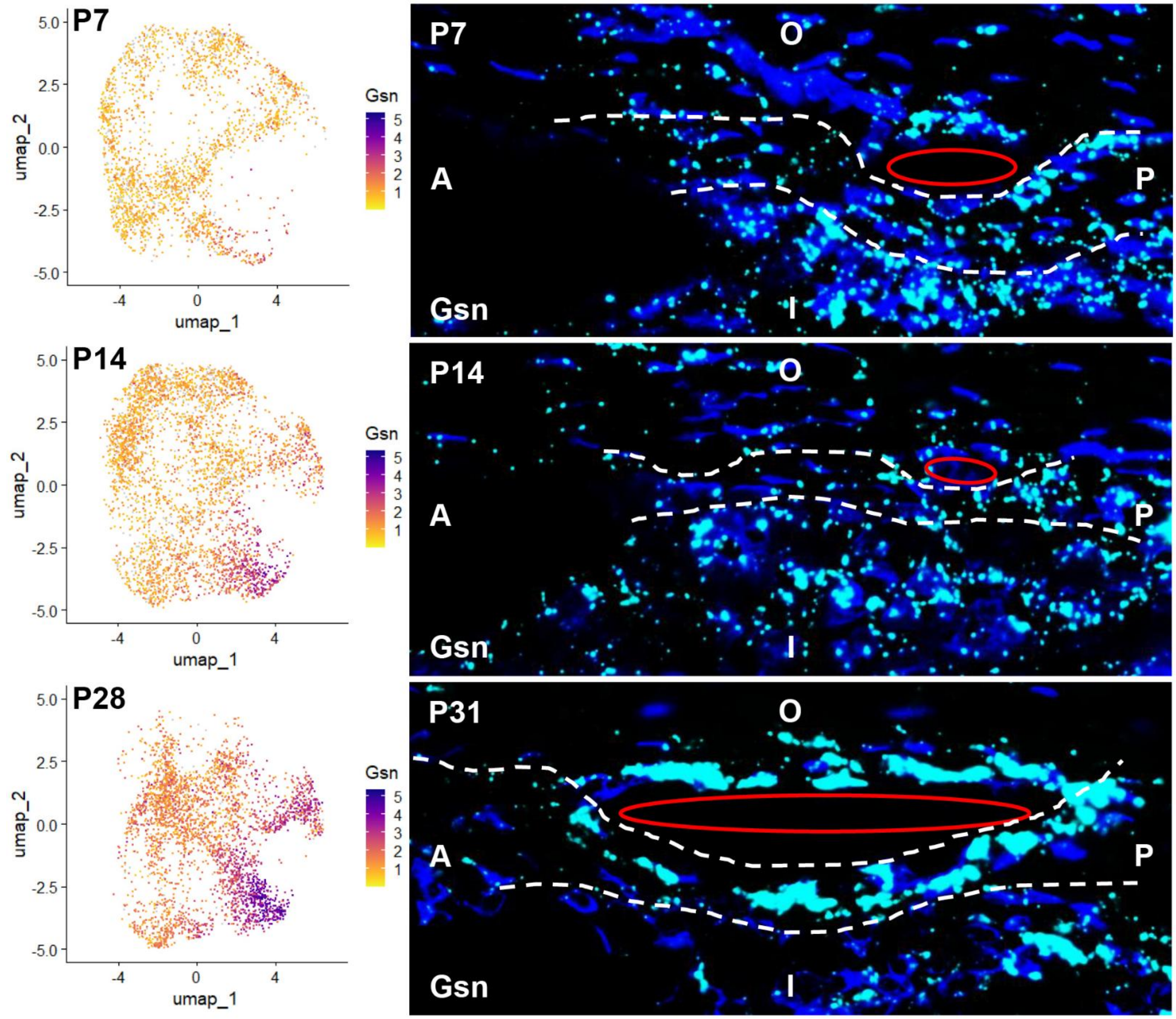


**Figure S13: Expression of *Gsn* within TMCs.**

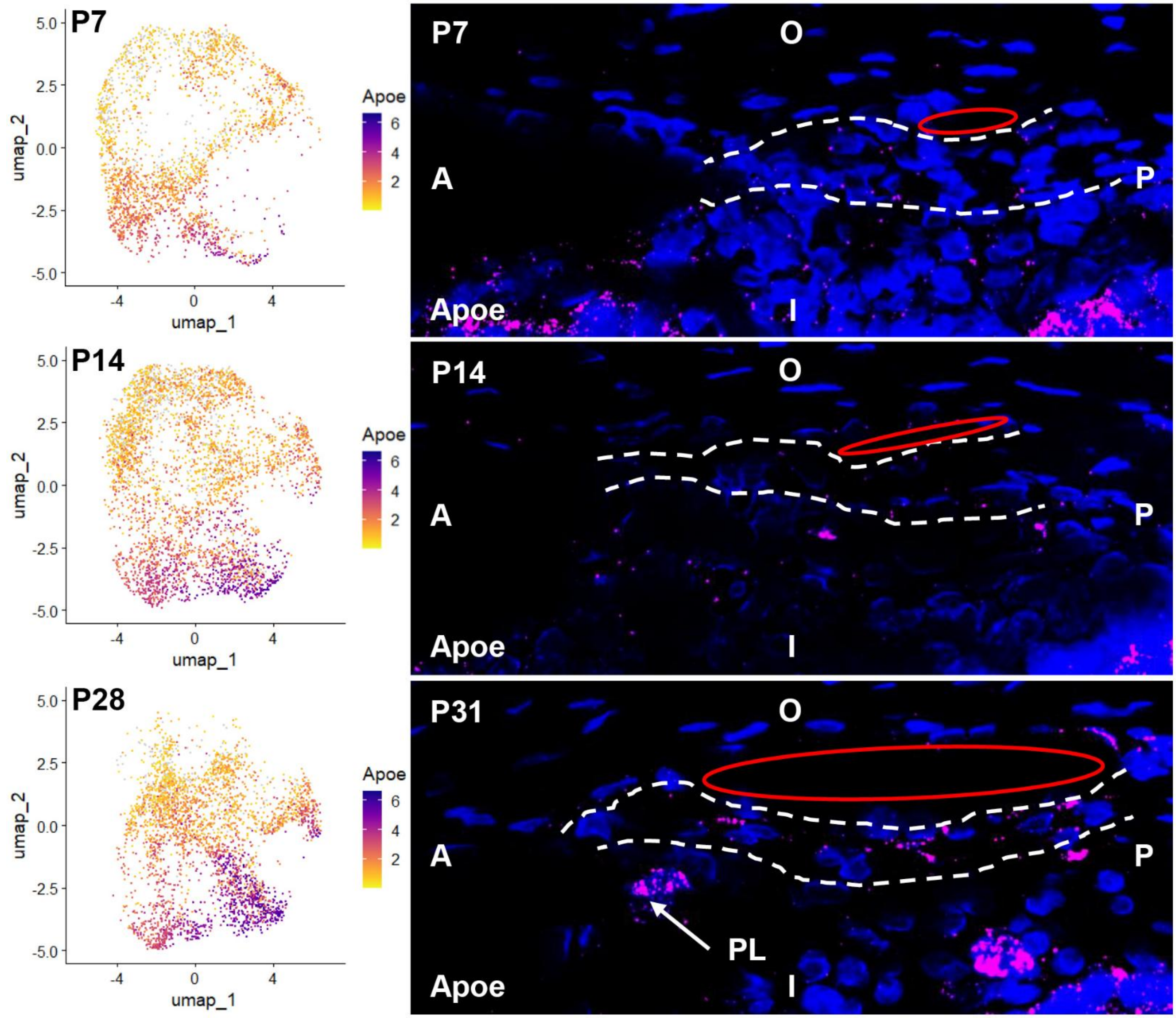


**Figure S14: Expression of *Apoe* within TMCs.**

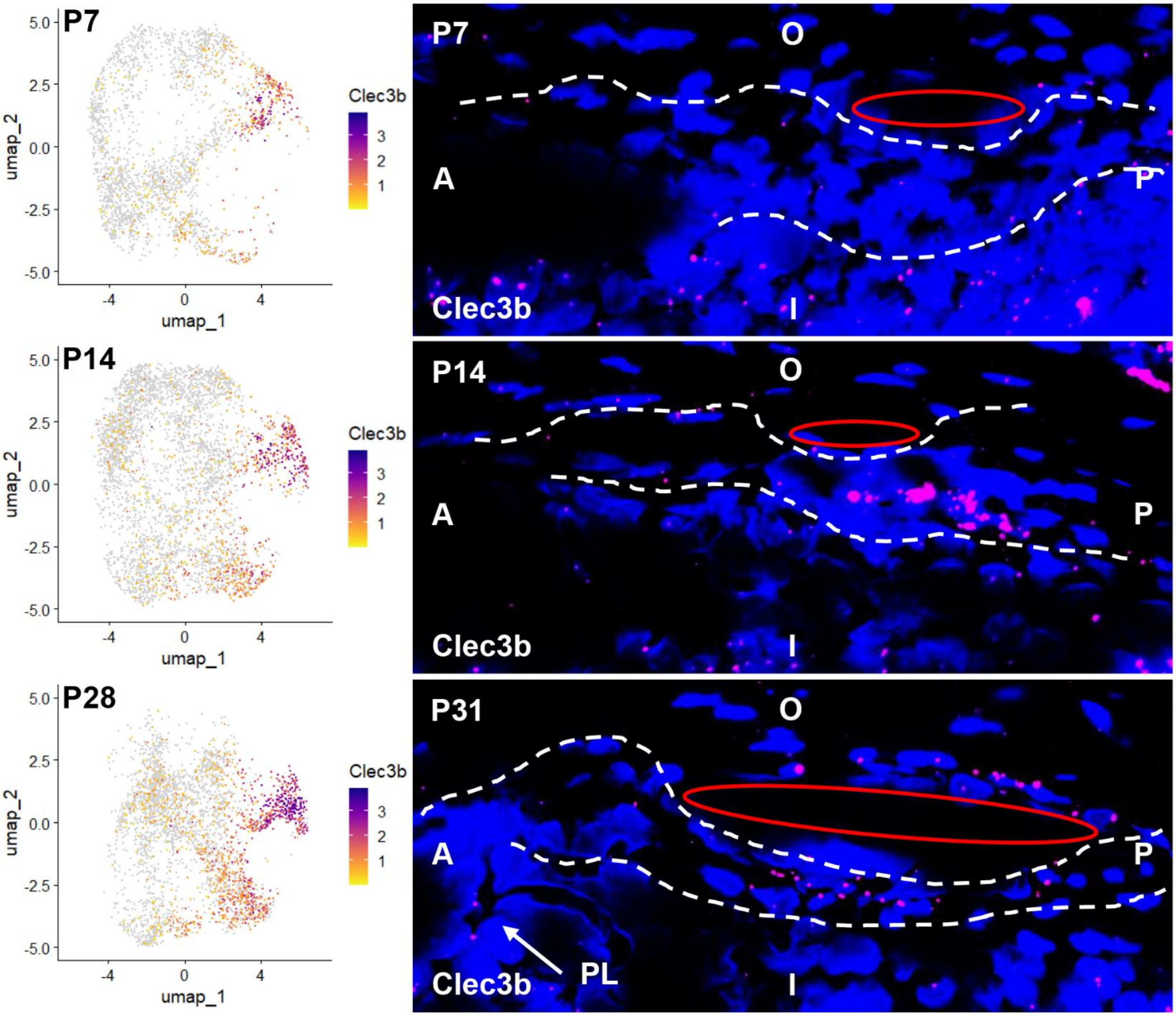


**Figure S15: Expression of *Clec3b* within TMCs.**

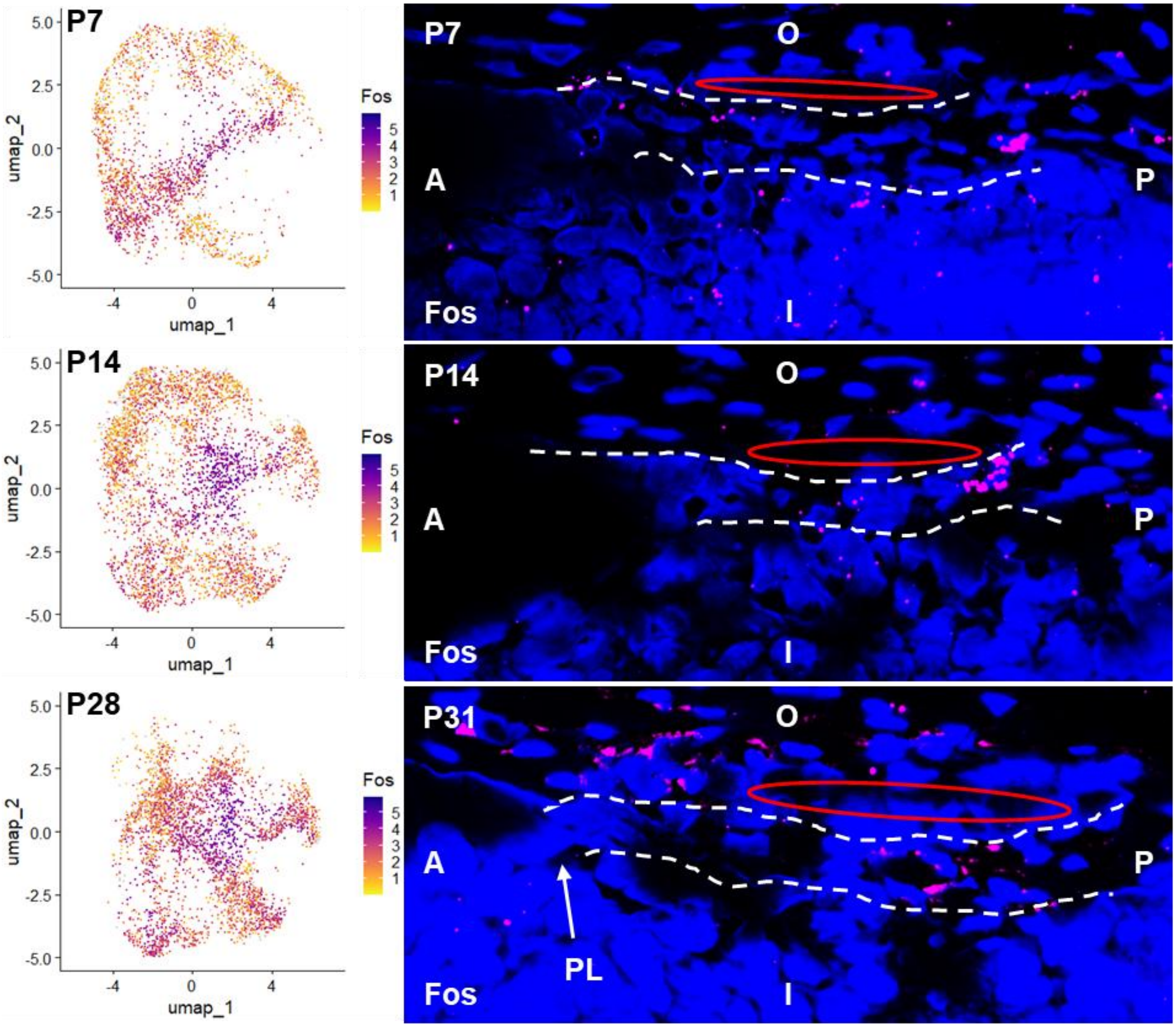


**Figure S16: Expression of *Fos* within TMCs.**

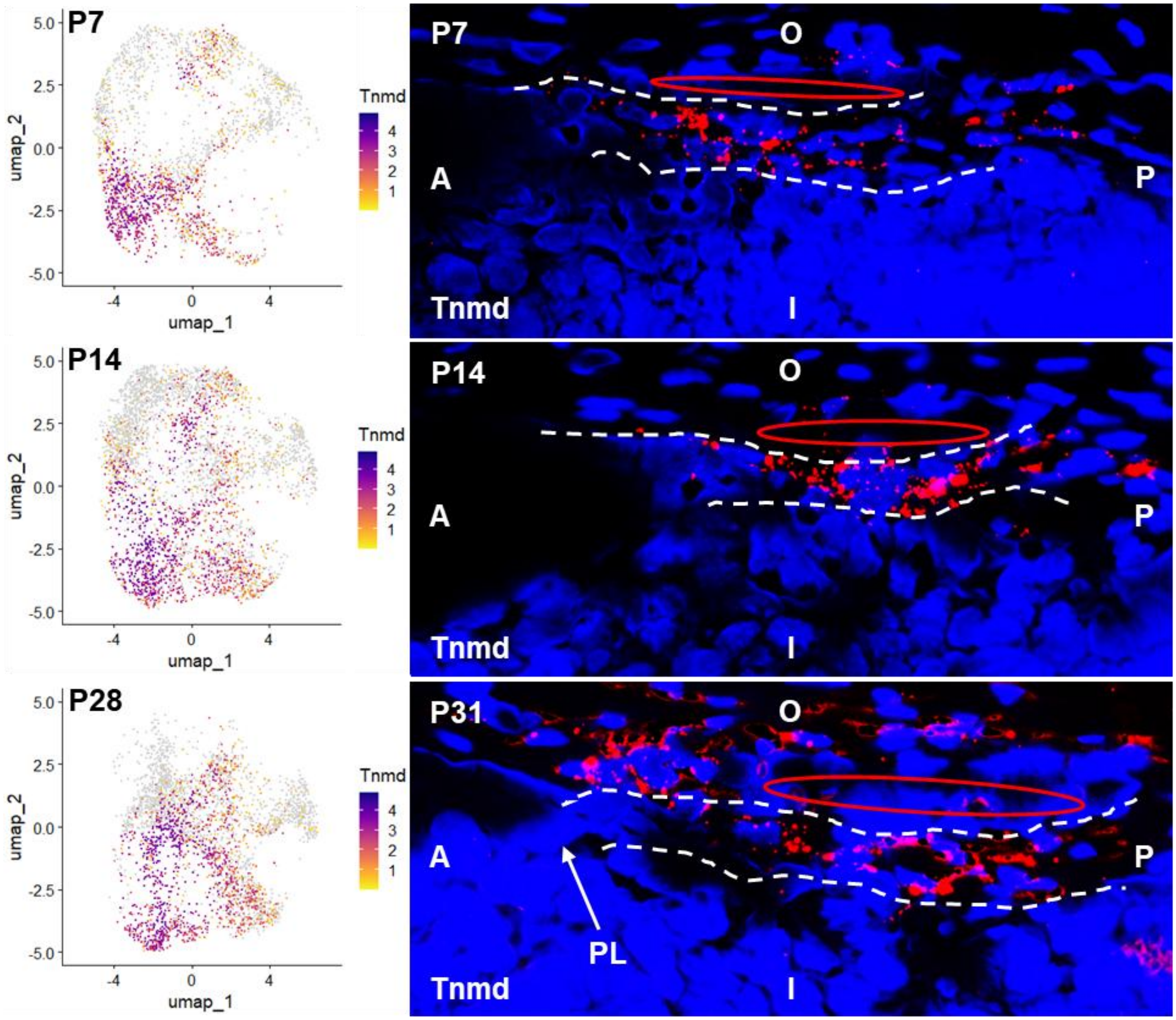


**Figure S17: Expression of *Tnmd* within TMCs.**

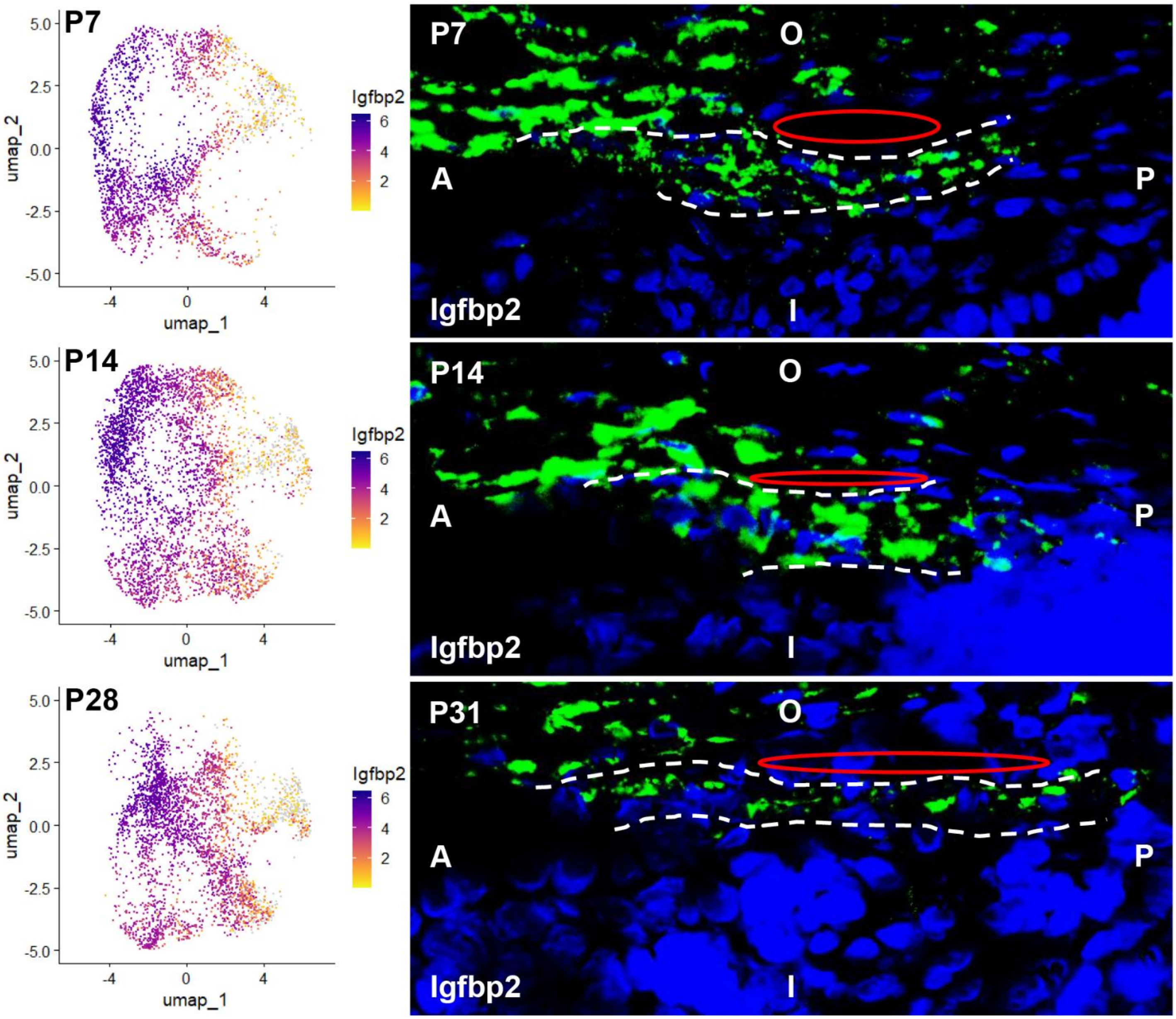


**Figure S18: Expression of *Igfbp2* within TMCs.**

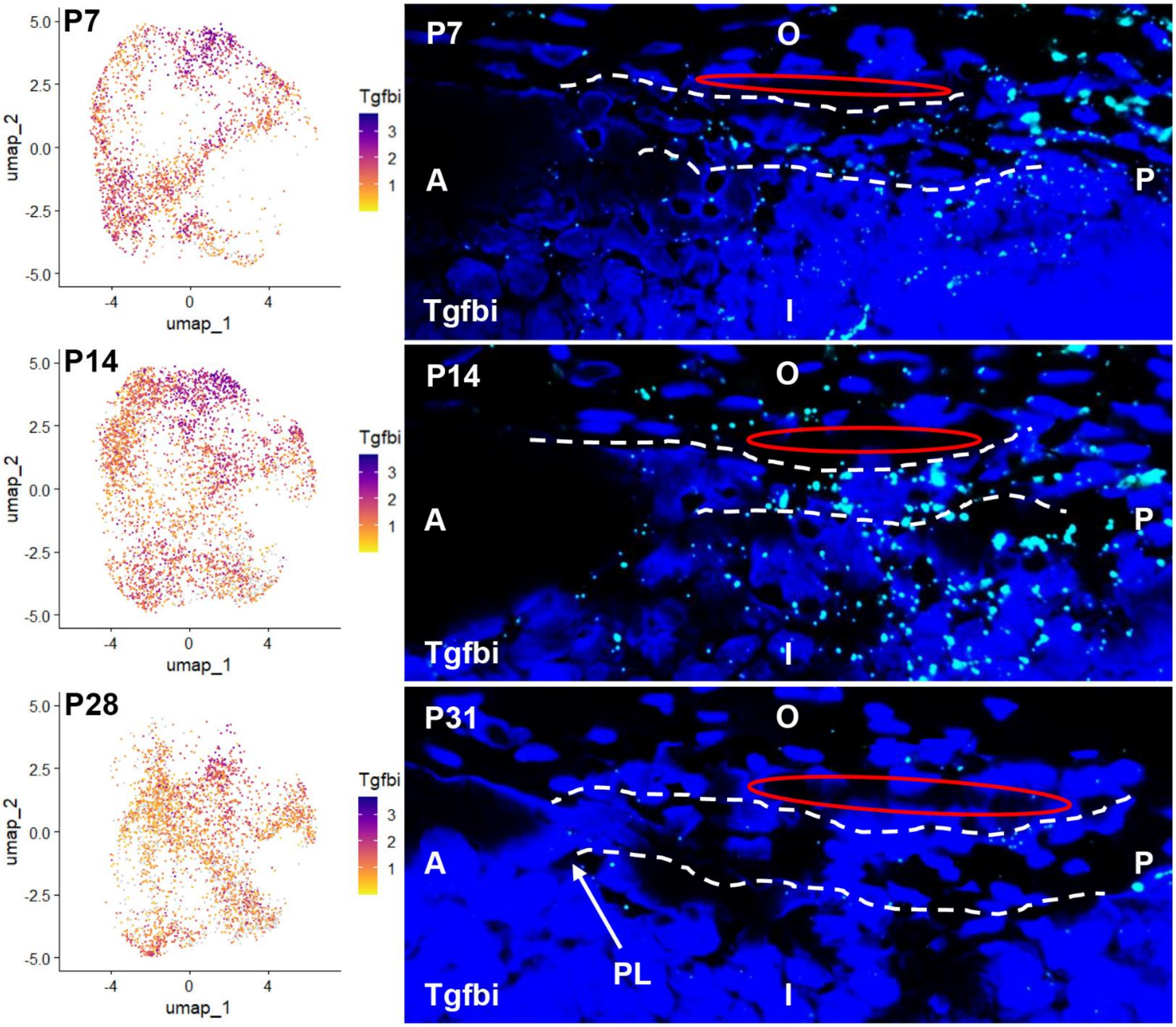


**Figure S19: Expression of *Tgfbi* within TMCs.**

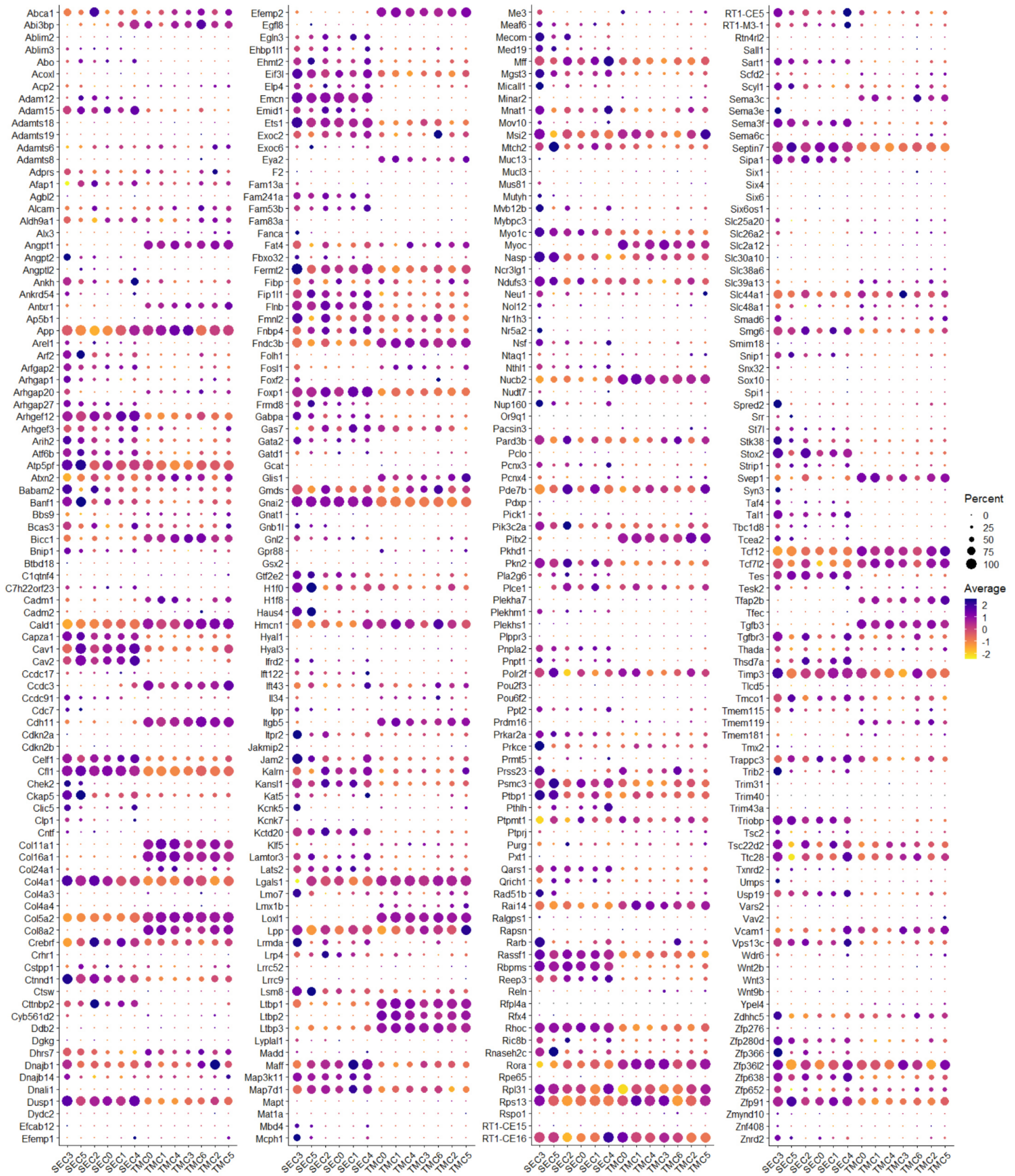


**Figure S20: Dot plot visualizing relative expression of 372 genes associated with IOP and POAG within SC- and TM-related cell type clusters.** Gene names listed on y-axis and cell type cluster names along x-axis; six cell clusters related to SC (SEC0-SEC5) and seven related to TM (TMC0-TMC6). Color denotes average relative gene expression normalized on a log2 scale. Dot size represents percentage of cells expressing the gene.

### Supporting Tables

**Table S1: Profiled cell numbers by major cell type cluster and tissue sample.** Columns show cell contributions from the 13 major cell type clusters (C0-C12) and rows provide contributions from individual limbal tissue strip samples. P, postnatal age; M, male; F, female; S, sample number; Epi, epithelial cells; Endo, endothelial cells; Prolif_Mes, proliferating mesenchymal cells.

| | C0 | C1 | C2 | C3 | C4 | C5 | C6 | C7 | C8 | C9 | C10 | C11 | C12 | Sum | Percent |
|---|---|---|---|---|---|---|---|---|---|---|---|---|---|---|---|
| **P7-M (S1)** | 1740 | 1129 | 515 | 405 | 444 | 145 | 67 | 105 | 109 | 139 | 25 | 116 | 14 | **4953** | **5.7** |
| **P7-F (S4)** | 1364 | 1279 | 324 | 304 | 310 | 126 | 92 | 81 | 128 | 96 | 26 | 74 | 12 | **4216** | **4.9** |
| **P8-M (S9)** | 2051 | 913 | 418 | 369 | 393 | 127 | 169 | 39 | 122 | 107 | 27 | 77 | 15 | **4827** | **5.6** |
| **P8-F (S12)** | 2018 | 1323 | 302 | 382 | 413 | 156 | 75 | 84 | 149 | 123 | 41 | 115 | 15 | **5196** | **6.0** |
| **P9-M (S14)** | 414 | 245 | 522 | 98 | 54 | 41 | 69 | 23 | 32 | 62 | 10 | 13 | 12 | **1595** | **1.8** |
| **P9-F (S17)** | 1504 | 1006 | 357 | 378 | 156 | 70 | 152 | 58 | 149 | 102 | 27 | 48 | 13 | **4020** | **4.6** |
| **P14-M (S18)** | 1541 | 1602 | 487 | 438 | 165 | 145 | 155 | 38 | 211 | 137 | 31 | 44 | 19 | **5013** | **5.8** |
| **P14-F (S15)** | 1408 | 1120 | 1049 | 412 | 360 | 216 | 317 | 59 | 217 | 230 | 40 | 46 | 24 | **5498** | **6.3** |
| **P15-M (S5)** | 1201 | 1651 | 240 | 399 | 224 | 146 | 104 | 196 | 129 | 165 | 32 | 35 | 11 | **4533** | **5.2** |
| **P15-F (S2)** | 1022 | 1129 | 339 | 382 | 241 | 171 | 112 | 278 | 126 | 158 | 23 | 36 | 15 | **4032** | **4.7** |
| **P16-M (S7)** | 909 | 1599 | 505 | 392 | 187 | 207 | 251 | 150 | 181 | 164 | 39 | 16 | 17 | **4617** | **5.3** |
| **P16-F (S10)** | 1270 | 1607 | 477 | 441 | 273 | 224 | 243 | 310 | 189 | 165 | 55 | 19 | 26 | **5299** | **6.1** |
| **P27-M (S3)** | 1854 | 1063 | 401 | 717 | 460 | 615 | 222 | 198 | 197 | 313 | 69 | 23 | 39 | **6171** | **7.1** |
| **P27-F (S6)** | 1182 | 1168 | 444 | 499 | 252 | 483 | 316 | 813 | 219 | 150 | 94 | 18 | 32 | **5670** | **6.5** |
| **P28-M (S11)** | 1204 | 1247 | 543 | 579 | 237 | 442 | 537 | 26 | 249 | 186 | 113 | 9 | 53 | **5425** | **6.3** |
| **P28-F (S8)** | 1118 | 1576 | 371 | 463 | 292 | 517 | 275 | 360 | 213 | 199 | 89 | 12 | 23 | **5508** | **6.4** |
| **P29-M (S13)** | 1011 | 978 | 671 | 479 | 175 | 390 | 466 | 145 | 316 | 214 | 83 | 5 | 34 | **4967** | **5.7** |
| **P29-F (S16)** | 881 | 1100 | 629 | 457 | 196 | 469 | 478 | 245 | 219 | 227 | 155 | 9 | 48 | **5113** | **5.9** |
| **Sum** | **23692** | **21735** | **8594** | **7594** | **4832** | **4690** | **4100** | **3208** | **3155** | **2937** | **979** | **715** | **422** | **86653** | **100** |
| **Percent** | **27.3** | **25.1** | **9.9** | **8.8** | **5.6** | **5.4** | **4.7** | **3.7** | **3.6** | **3.4** | **1.1** | **0.8** | **0.5** | **100** | **0** |

**Table S2: Tissue acquisition groupings for 18 limbal tissues acquired over a 3-day period.** The six groups of three tissue samples comprised a systematic mix of developmental ages (P7-P29) and genders (M or F) to limit acquisition bias. Each day, a group was processed in the morning (AM) and another group in the afternoon (PM).

| Group | Acquisition Time | Sample-1 | Sample-2 | Sample-3 |
|---|---|---|---|---|
| 1 | Day-1 AM | P7-M | P15-F | P27-M |
| 2 | Day-1 PM | P7-F | P15-M | P27-F |
| 3 | Day-2 AM | P16-M | P28-F | P8-M |
| 4 | Day-2 PM | P16-F | P28-M | P8-F |
| 5 | Day-3 AM | P29-M | P9-M | P14-F |
| 6 | Day-3 PM | P29-F | P9-F | P14-M |

**Table S3: Rat probes used in ISH assays (RNAscope).** Probes were designed and synthesized by Bio-Techne (https://www.bio-techne.com/reagents/rnascope-ish-technology). Probe channels shown as C1-C4 (C1 where channel not stated). Opal dyes were diluted at 1:1000 or 1:1500.

| Probe | Cat # | Fluorophore | Dye Dilution |
|---|---|---|---|
| Rn-Angpt1 | 837021 | Opal 570 | 1:1000 |
| Rn-Apoe-C3 | 1690251-C3 | Opal 570 | 1:1500 |
| Rn-Ccl21 | 878391 | Opal 570 | 1:1000 |
| Rn-Cd34-C2 | 1055971-C2 | Opal 650 | 1:1500 |
| Rn-Chad | 890651 | Opal 650 | 1:1000 |
| Rn-Clec3b-C4 | 1809361-C4 | Opal 570 | 1:1000 |
| Rn-Cyp1b1-C4 | 493211-C4 | Opal 570 | 1:1000 |
| Rn-Flt1-C4 | 435701-C4 | Opal 650 | 1:1000 |
| Rn-Fos | 403591 | Opal 570 | 1:1500 |
| Rn-Gjc2-C2 | 1598241-C2 | Opal 650 | 1:1000 |
| Rn-Gsn-C3 | 828901-C3 | Opal 650 | 1:1000 |
| Rn-Igfbp2-C1 | 1171961-C1 | Opal 520 | 1:1500 |
| Rn-Ltbp2-C3 | 1589761-C3 | Opal 570 | 1:1000 |
| Rn-Myoc-C2 | 814801-C2 | Opal 520 | 1:1500 |
| Rn-Nox4 | 452791 | Opal 570 | 1:1000 |
| Rn-Pecam1-C3 | 315311-C3 | Opal 650 | 1:1500 |
| Rn-Pitx2 | 543091 | Opal 650 | 1:1000 |
| Rn-Prox1-C1 | 1088941-C1 | Opal 570 | 1:1000 |
| Rn-Sele-C4 | 1809371-C4 | Opal 650 | 1:1500 |
| Rn-Svep1-C1 | 1598301-C1 | Opal 650 | 1:1000 |
| Rn-Tek-C2 | 435721-C2 | Opal 570 | 1:1000 |
| Rn-Tgfbi-C4 | 888731-C4 | Opal 650 | 1:1500 |
| Rn-Tnmd-C3 | 436931-C3 | Opal 620 | 1:1500 |
| Rn-Top2a-C2 | 1097631-C2 | Opal 570 | 1:1500 |
